# Interactive visualisation of raw nanopore signal data with Squigualiser

**DOI:** 10.1101/2024.02.19.581111

**Authors:** Hiruna Samarakoon, Kisaru Liyanage, James M. Ferguson, Sri Parameswaran, Hasindu Gamaarachchi, Ira W. Deveson

**Author notes:** Joint senior authors; correspondence.

## Abstract

Nanopore sequencing measures ionic current during the translocation of DNA, RNA or protein molecules through a nanoscale protein pore. This raw current signal data can be ‘basecalled’ into sequence information and has the potential to identify other diverse molecular features, such as base modifications, secondary structures, etc. Despite the unique properties and potential utility of nanopore signal data, there are currently limited options available for signal data visualisation. To address this, we have developed *Squigualiser*, a toolkit for intuitive, interactive visualisation of sequence-aligned signal data, which currently supports both DNA and RNA sequencing data from Oxford Nanopore Technologies (ONT) instruments. A series of methodological innovations enable efficient alignment of raw signal data to a reference genome/transcriptome with single-base resolution. *Squigualiser* generates an interactive signal browser view (HTML file), in which the user can navigate across a genome/transcriptome region and customise the display. Multiple independent reads are integrated into a signal ‘pileup’ format and different datasets can be displayed as parallel tracks to facilitate their comparison. *Squigualiser* provides the most sophisticated framework for nanopore signal data visualisation to date and will catalyse new advances in signal analysis. We provide *Squigualiser* as an open-source tool for the nanopore community: https://github.com/hiruna72/squigualiser

## INTRODUCTION

Nanopore sequencing enables direct analysis of native DNA, RNA and protein molecules, with countless potential applications across the life sciences^1^. Devices from Oxford Nanopore Technologies (ONT) measure the displacement of ionic current as a DNA or RNA molecule passes through a nanoscale protein pore. The device records time-series current signal data – commonly referred to as ‘squiggle’ data – which can be ‘basecalled’ into sequence reads and/or analysed directly^1^.

This electrophysical sequencing mechanism sets nanopore sequencing apart from alternative platforms. Other existing high-throughput sequencing technologies rely on optical detection of fluorescently labelled DNA nucleotides^2,3^ – a discrete sequencing mechanism. In contrast, nanopore sequencing generates continuous data in both the time and amplitude domains, and spanning a wide dynamic range. The data is shaped by any molecular feature that influences the flow of ionic current through a pore occupied by a DNA/RNA molecule and/or the rate at which the molecule moves through the pore (the translocation speed). As a result, nanopore sequencing is amenable to detection of diverse ‘modified’ DNA^4,5^ or RNA^6^ nucleotides, DNA damage^7^, RNA secondary structures^8,9^, or other features beyond the primary nucleotide sequence. Early stage demonstrations of protein sequencing further show the suitability of nanopore signal data for determining amino acid sequence and post-translational modifications^10^.

However, nanopore signal data is large, complex and unfamiliar to many in the genomics community. Despite success in detection of some DNA and RNA modifications, most notably 5-methylcytosine (5mC)^4^, these challenges have, arguably, constrained the development of signal-level analysis methods, with most users preferring to work exclusively with basecalled sequence data.

One key barrier is a lack of effective methods for visualisation of nanopore signal data. To address this, we have developed *Squigualiser* (<u>**Squig**</u>gle vis<u>**ualiser**</u>), a new framework for intuitive signal data visualisation. *Squigualiser* builds upon existing methodology for signal-to-sequence alignment in order to anchor raw signal data points to their corresponding positions within basecalled reads or within a reference genome/transcriptome sequence. A new encoding technique (the *ss tag*) enables efficient, flexible representation of signal alignments and normalises outputs from alternative alignment tools. A new method for k-mer-to-base shift correction addresses ambiguity in signal alignments to enable visualisation of genetic variants, modified bases, or other features, at single-base resolution. The *Squigualiser* toolkit uses these and other underlying innovations to process raw or pre-aligned signal data and generate an interactive browser view for data exploration.

## RESULTS

### Sequence-aligned signal data visualisation

*Squigualiser* is a toolkit for visualising nanopore signal data, which currently supports both DNA and RNA sequencing data from ONT instruments. *Squigualiser* provides the necessary tools to integrate raw signal data, basecalled reads, a reference genome/transcriptome sequence and other genomic feature files, into an intuitive visual format. The software generates an interactive browser view (HTML format) capturing a specified set of reads and/or reference region, which can be displayed and navigated in a modern web browser. A pre-built example is provided in **Supplementary_File_1.html** and a brief instructional video is available at the following link: https://youtu.be/kClYH4KpOjk.

To provide the molecular context for signal interpretation, *Squigualiser* displays raw signal data points in alignment with their corresponding nucleotides within a basecalled read (signal-to-read) or a reference genome/transcriptome sequence (signal-to-reference). Values for an individual nanopore read are plotted as a string of consecutive points against a vertical axis specifying the measured current (in pico-amps) and a horizontal axis representing the passage of time as the molecule was sequenced (**Fig1a**). The horizontal axis is partitioned and coloured according to the underlying bases (A,C,G,T[U]) in the read/reference sequence. Because translocation speed is not constant, the number of signal data points per nucleotide varies within and between reads. To account for this, when viewing signal-to-read data, the horizontal axis may be alternatively displayed such that the distance between consecutive signal values is equal (time-scale) or dynamically re-packed such that each nucleotide has equal width (molecule-scale; **Fig1a**).

**Fig1.**
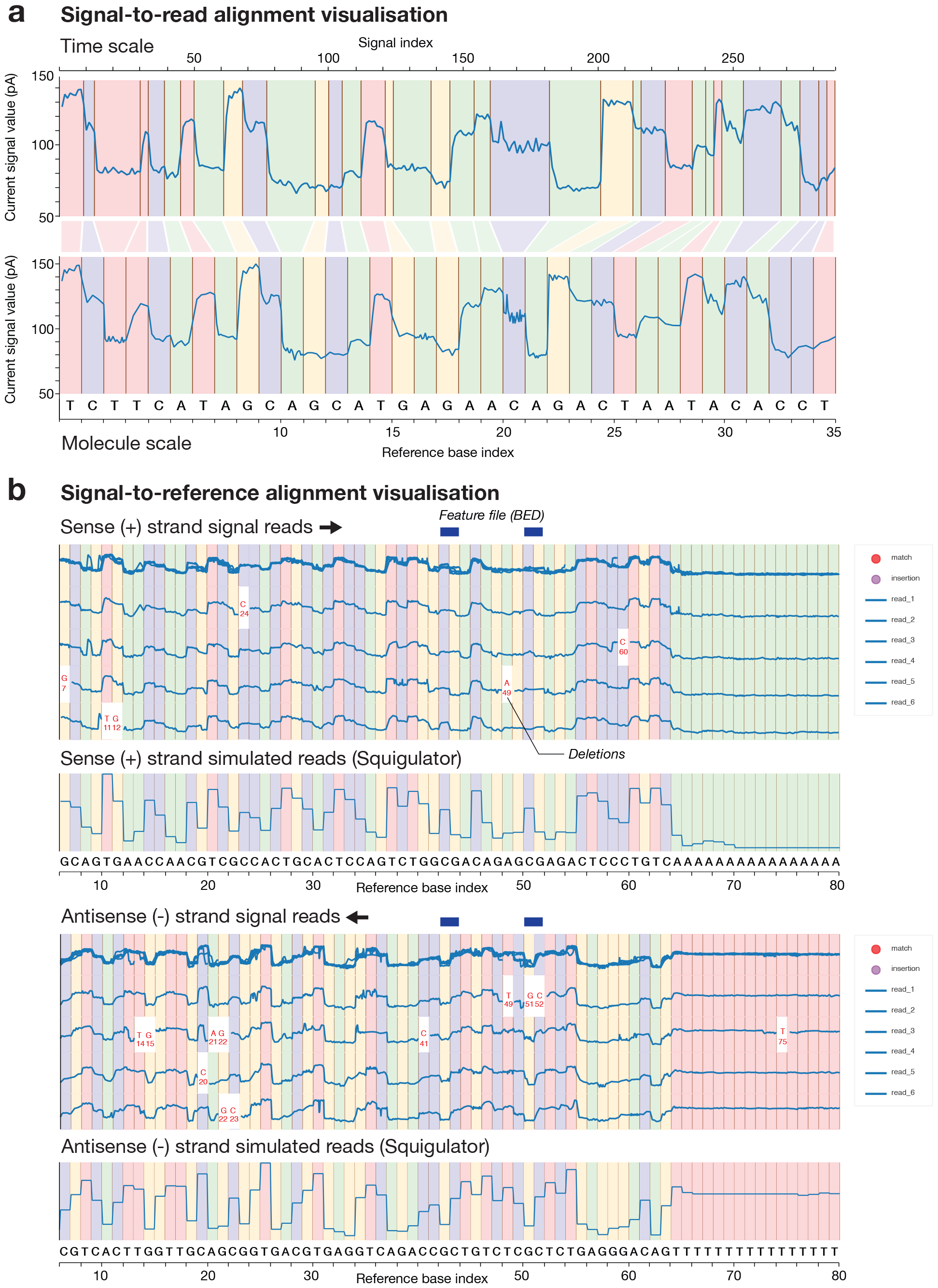
Visualising sequence-aligned nanopore signal data with *Squigualiser*. (**a**) Exported images from *Squigualiser* signal viewer show signal-to-read alignments for a ∼35 nucleotide subsequence within a single ONT read. In the upper image, aligned signal data is displayed in ‘time-scale’, meaning signal values are spaced uniformly. In the lower image, the same data is displayed in ‘molecule-scale’, wherein signal values are dynamically re-packed such that their aligned nucleotides are displayed with equal width, regardless of how many signal values are aligned to a given base. (**b**) Exported images show signal-to-reference alignments for a ∼80 nucleotide reference genome region, highlighting various features of the *Squigualiser* visualisation strategy. By using molecule-scale, multiple independent reads can be overlaid in a consistent pileup despite variable translocation speeds within and between reads. Reads may be selected / deselected using the control panel (right), to determine which are actively displayed in the pileup. Reads aligned to the antisense (-) strand in DNA sequencing data are shown separately, and in reverse signal orientation (reverse-complement molecule-scale) so that complementary sense/antisense bases are aligned. Simulated signal data from *Squigulator* may be displayed as a parallel track/s to assist signal interpretation.

Signal-to-reference alignment enables a user to inspect data from multiple independent reads and/or datasets corresponding to the same genome or transcriptome locus. To facilitate this, *Squigualiser* provides a signal ‘pileup’ view, where individual reads are stacked vertically, aligned in molecule-scale with the underlying reference sequence, and all reads are overlaid in a single track above (**Fig1b**). The user can interactively select/deselect reads to determine which are actively displayed within the pileup. For DNA sequencing data, signal alignments to the antisense (-) strand are displayed in reverse orientation, so that data points are aligned with their corresponding base in the genome reference, and sense (+) and antisense (-) reads are organised in separate pileup tracks (**Fig1b**). Multiple independent datasets may also be displayed as separate, parallel tracks, similar to the layout in popular genome browsers, such as Integrative Genomics Viewer (IGV)^11^. A coordinate annotation file (BED format) may be displayed, in order to identify relevant features within the reference (e.g. methylated CpG sites; **Fig1b**).

Our recent tool for nanopore signal data simulation, *Squigulator*^*12*^, can be used to generate and display a simulated data track within *Squigualiser*, based on the underlying read/reference sequence. This ‘expected’ data track provides useful context, with deviations in the accompanying experimental data being potentially indicative of sequence variants, modified bases or other features (**Fig1b**). Insertions and deletions commonly occur within signal-to-reference alignments, due to a combination of basecalling and/or alignment errors and true genetic variants. To accommodate this, *Squigualiser* leaves empty space within aligned signal reads at the positions of deleted nucleotides and labels the identity of missing bases (**Fig1b**). Signal data points corresponding to inserted bases are stacked into a single point at the border of the preceding reference nucleotide and can be revealed with the cursor.

### Signal alignment methods

Signal-to-sequence alignment is central to this visualisation framework and several methodological innovations were required to streamline and optimise this process. *Squigualiser* reads and writes signal alignments in BAM and PAF formats familiar to the genomics community, but customised for signal data. For example, we developed a new method for compressed encoding of signal alignments within BAM or PAF format, an auxiliary tag named the ‘*ss tag*’, which is described in detail in **Supplementary Note 1**. Inspired by the CIGAR string from sequence alignment format^13^, the *ss tag* encodes signal-to-sequence matches, insertions or deletions of varying lengths in a minimal alphanumeric string. Measured on a typical dataset, this has a superior compression ratio (∼1.6) than the ‘move table’ format used by ONT basecalling software (*Guppy*/*Dorado*), among other advantages.

Perhaps more importantly, the *ss tag* serves to normalise the signal alignment representations used by different software. This allows *Squigualiser* to support alignment data generated through a variety of alternative methods. The simplest approach is to directly extract and re-format signal-to-read alignment information from the ONT move table generated during basecalling, using the *Squigualiser reformat* subtool (see **Supplementary Note 2**; see **Methods**). This may also be integrated with basecalled read alignments (BAM format) from *minimap2*^*14*^ or another standard aligner, in order to generate signal-to-reference alignments. This is done using the *Squigualiser realign* subtool (see **Methods**).

Alternatively, the user may opt to use various external signal alignment software, including *F5c resquiggle*^*15*^ for signal-to-read alignment, or *F5c eventalign*^*15*^, *Nanopolish*^*4*^ signal projection, or *Sigfish*^*16*^ dynamic time warping methods for signal-to-reference alignment (**Fig2a**; see **Methods**). Each approach has different merits and the user is free to choose their preferred method, depending on the context. Moreover, in supporting different methods, *Squigualiser* provides a framework to compare and evaluate them. For example, when assessed by the precision and uniformity of signal alignments across multiple independent reads, *F5c eventalign* achieves superior performance and is generally our preferred approach (**Fig2a,b**). Although this analysis shows the *reformat* / *realign* method to be the least precise, this method has the advantage of being non-reliant on a k-mer pore model or external alignment software (see **Methods**).

**Fig2.**
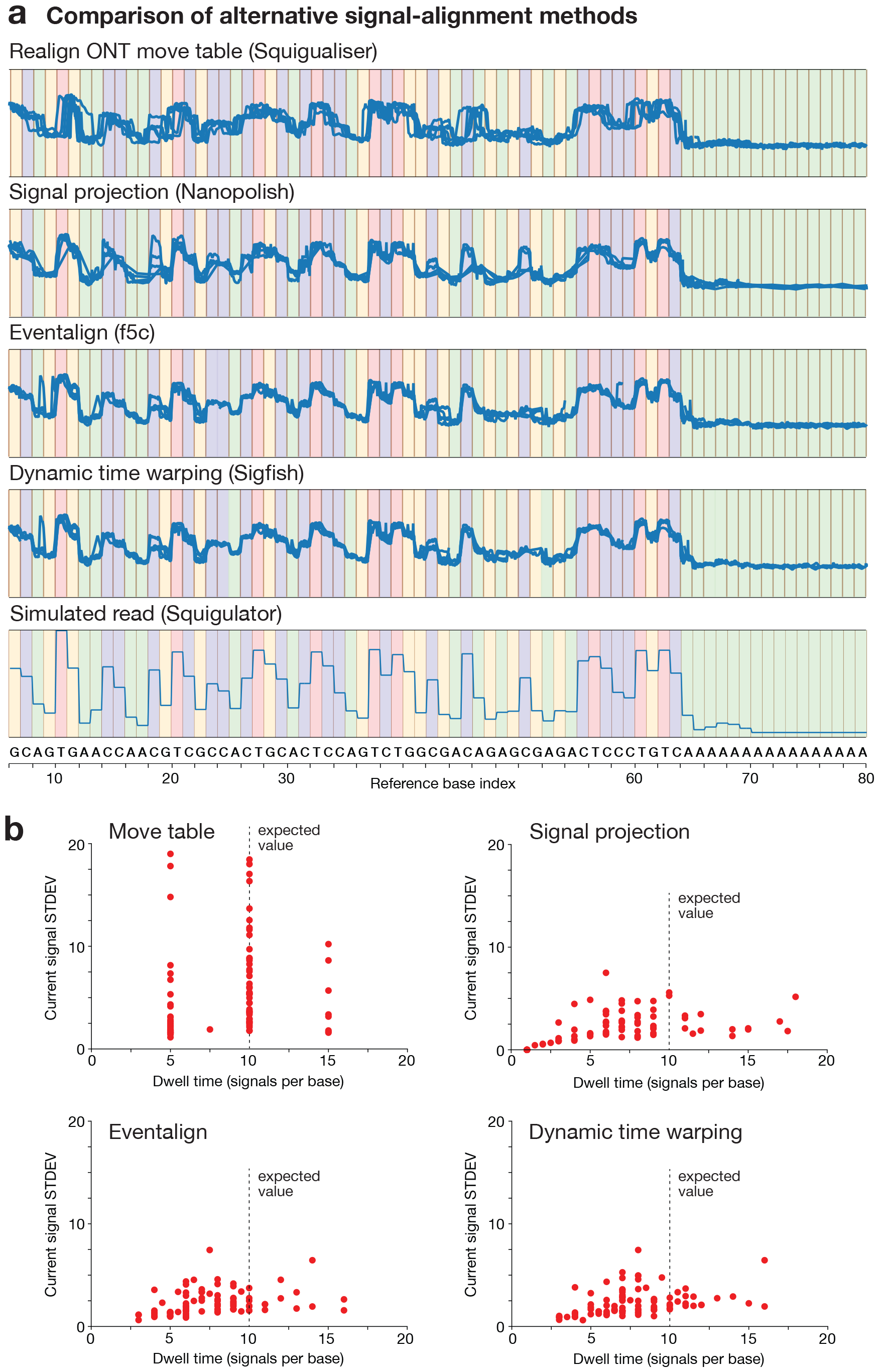
Using *Squigualiser* to evaluate alternative signal alignment methods. (**a**) Exported *Squigualiser* pileup images show a matched set of signal-to-reference alignments generated using four alternative methods: alignments extracted directly from the ONT move table using *Squigualiser reform* and *realign* subtools; alignments generated using the signal projection method in the *Nanopolish* toolkit; alignments generated using the event alignment method in the *F5c* toolkit; alignments generated using *Sigfish* dynamic time warping for signal mapping. The bottom track shows simulated signal data from *Squigulator*, as a guide. (**b**) The alternative methods above can be statistically evaluated on the precision/uniformity of signal alignments. Dots plots here show standard deviations for all signal data values aligned to each individual reference base (each base is a single red dot) on the vertical axis and the number of signal values per base per read on the horizontal axis. *F5c* eventalign shows the lowest variation across the two domains, and is generally our preferred method for signal-to-reference alignment.

### K-mer-to-base shift correction

One impediment to intuitive visualisation is the lack of resolution during signal alignment. Existing signal-to-read and signal-to-reference alignment methods work by anchoring ‘events’ (step changes in signal values) not to individual nucleotides but to k-mers (DNA/RNA sub-sequences of length = k). Therefore, there is an arbitrary offset (distance n; where n < k) between a given base and the most appropriate event to represent that base in the aligned signal. This makes it difficult to precisely associate signal events with genetic variants, DNA/RNA modifications, or other features at single-base resolution.

It is known that specific bases have predictable effects on the current level during nanopore sequencing, as determined by their size, shape, charge, etc^17^. In ONT data, for example, T nucleotides are associated with the highest current. We reasoned that these effects could be used to calculate and correct the arbitrary offset in signal alignments, for more intuitive visualisation. To do so, we identify the base that is predicted to be most significant within a given k-mer (i.e. the single base with the greatest influence on the current level), infer the local signal event with which it is likely associated, and measure the offset between them. Applied globally to a given dataset, this identifies an offset value by which all signal alignments in the dataset should be shifted in order to align all bases with their most relevant signal events. This is protocol-specific. For example, on DNA sequencing data generated from an R10.4.1 flow-cell at 400bps translocation speed, basecalled with *Guppy* SUP model (dna_r10.4.1_e8.2_sup@v3.5.1) and aligned with *F5c eventalign*, we find a consistent offset of -6bp between the most significant base within a given k-mer and its corresponding event (**Fig3a**). However, the appropriate offset value differs for different flow-cell versions, basecalling models and/or signal alignment methods, and therefore needs to be re-calculated for a given protocol. *Squigualiser* includes a subtool *calculate_offsets* to identify the appropriate base-shift for a given dataset, in addition to pre-calculated offsets for many common protocols/workflows (see **Methods**). This k-mer-to-base shift correction strategy is described in detail in **Supplementary Note 3**.

**Fig3.**
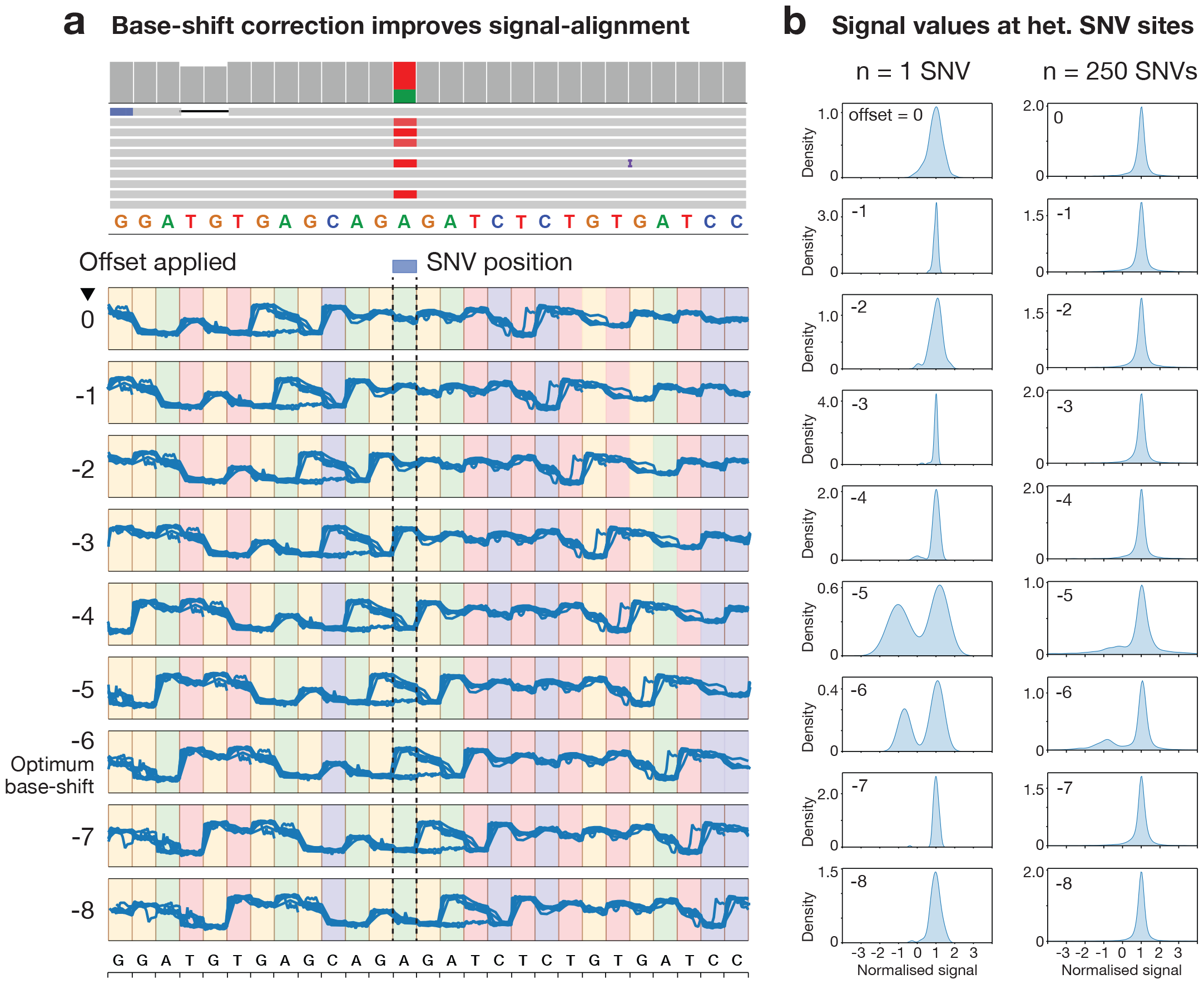
Evaluating *Squigualiser* k-mer-to-base shift correction. The k-mer-to-base shift correction strategy is designed to optimise signal alignment visualisation by correcting the arbitrary offset between a given nucleotide and its most relevant corresponding event in aligned signal. *Squigualiser* calculates and corrects this offset for a given protocol/dataset; for the dataset displayed here, an offset of -6 was determined by *Squigulator* to be optimum. To test whether this correction led to improved resolution, we inspected known heterozygous A-to-T SNVs, reasoning that these sites should show bimodal signal values corresponding to the reference and alternate alleles, and that the optimum offset should show the highest degree of bimodality at these sites. (**a**) The upper track shows basecalled read alignments to a single example SNV displayed in the IGV genome browser. The lower tracks show signal-alignment pileup images from *Squigualiser* for the same set of reads, and the same SNV site, with a range of different offset values applied (0 to -8). (**b**) Histograms show the density of signal values at a single SNV site (same site as in **a**) and across 250 randomly selected SNV sites in the same dataset. For both the single SNV and the full set, the optimum offset of -6 calculated by *Squigualiser* is associated with the highest degree of bimodality.

To evaluate our new k-mer-to-base shift correction strategy, we extracted all signal values aligned to the site of a known heterozygous SNV within the HG002 genome reference sample, repeating this for a set of 250 A-to-T SNVs on chr22 across a range of possible offset values (0 to -8). We observed the highest degree of bimodality among signal values aligned to heterozygous SNVs when applying a k-mer-to-base shift correction of -6, which was also the optimum offset value calculated by *Squigualiser* for this protocol (**Fig3a,b**). Without any correction (offset = 0), signal values appeared unimodal at heterozygous SNV sites, thereby failing to capture expected current differences between the reference and alternate bases, which are typically visible at a nearby position (**Fig3a,b**). This analysis demonstrates how our k-mer-to-base shift correction approach leads to improved resolution in signal-alignment visualisation.

### Example use case: visualising DNA and RNA modifications

There are many foreseeable use-cases for the signal data visualisation frameworks provided by *Squigualiser*. To provide two simple examples, we used *Squigualiser* to visually inspect a set of modified DNA and RNA bases in ONT datasets (see **Supplementary Notes 4 & 5**). Modified DNA or RNA bases, such as the epigenetic regulator 5-methylcytosine (5mC), are associated with distortions of ONT signal data relative to their corresponding canonical bases, which are the basis for their detection. However, modifications are diverse, subtle and may be challenging to detect^18^. Given the capacity to visualise multiple independent signal reads and/or datasets (e.g. case vs control), aligned to their appropriate position in the reference genome/transcriptome with single-base resolution, *Squigualiser* is well-suited for visual inspection of candidate modifications.

To visualise 5mC DNA modifications, we processed a recent DNA sequencing dataset (R.10.4.1, LSK114) from the genome reference sample HG002 with *Squigualiser*. The dataset was basecalled (*Guppy*), aligned to the human reference genome (hg38; *minimap2*), and 5mC modifications were detected with *F5c call-methylation*. In parallel, we performed signal alignment with *F5c eventalign* and generated signal-to-reference data plots with *Squigualiser*. To illustrate how *Squigualiser* can be used to interrogate the underlying data, we located CpG sites detected as partially methylated, then partitioned reads predicted by *F5c* to be methylated vs unmethylated and displayed these in separate pileup tracks (see **Supplementary Note 4, Figure 1**). Visual inspection highlights the distinct signal pattern between methylated vs unmethylated reads aligned to CpG sites, assisting the user to validate/interpret the meth-calling results from *F5c*. This example is outlined in detail in **Supplementary Note 4**.

To visualise pseudouridine (Ψ) RNA modifications, we downloaded publicly available direct RNA sequencing data generated from synthetic RNA molecules produced with (Ψ+) and without (Ψ-) modifications in place of canonical U bases^19^. The dataset was basecalled (*Guppy*), aligned to the relevant synthetic reference sequences (*minimap2*), then signal alignments were generated with *F5c eventalign* and visualised with *Squigualiser*. Visual inspection showed poor alignment quality for the Ψ+ data, with many gaps appearing in regions where T(U) bases appear in the reference sequence. By retrieving signal values at these sites, we were able to create a customised ONT k-mer pore model that includes expected signal values for RNA k-mers containing Ψ modifications, and providing this pore model to *F5c eventalign* led to major improvements in signal alignment quality, with many fewer gaps observed in the Ψ+ data. This approach enabled visual comparison of aligned signal values at known Ψ+ vs Ψ-sites (see **Figure 7** within **Supplementary Note 5**), assisting the user to identify the effects of this RNA modification on the underlying signal. This example is outlined in detail in **Supplementary Note 5**.

## DISCUSSION

*Squigualiser* is a toolkit for visualisation of sequence-aligned nanopore signal data. *Squigualiser’s* signal-to-reference alignment visualisation is particularly useful, as it enables inspection of multiple independent reads and/or datasets anchored to their corresponding genome or transcriptome position. This provides the context to interpret signal data in relation to DNA variants, DNA/RNA modifications, or other molecular features impacting signal data. *Squgualiser’s* visualisation strategy depends on a series of methodological innovations enabling reliable and efficient signal alignment at single-base resolution, which are explored in detail in **Supplementary Notes 1-3**. The alignment and visualisation workflow is flexible, supporting multiple different alignment strategies and other customisable features.

While *Squigualiser* is not the first software for ONT signal visualisation, it constitutes a major advance in this area. For example, signal analysis toolkits *Poretools*^*20*^, *PoRe*^*21*^ and *Tombo* (https://github.com/nanoporetech/tombo) have sub-tools to support signal visualisation, however, these generate static plots with relatively crude reference alignment, and multiple data tracks cannot be displayed in parallel. *Uncalled4*^*22*^ can be used to generate a semi-interactive signal data plot in HTML format, which partly inspired our visualisation strategy for *Squigualiser*. However, *Uncalled4* is only has capacity to plot a single read in time-scaled signal-to-read format, which limits its utility for data exploration. Another standalone package for signal visualisation, *Bulkvis*^*23*^, is effective for visualising raw bulk-FAST5 files, but does not support signal-to-reference alignment, nor can multiple reads or datasets be displayed in parallel.

*Squigualiser* is the first interactive visualisation framework, generating a customisable browser view (HTML format), in which the user may navigate across a region, zoom in/out, select/deselect reads/tracks on display, and customise visualisation parameters. Dynamic packing of signal data points into their corresponding reference base, scaled by nucleotide rather than time (to account for variable translocation speed), and reversal of signal reads aligned to the DNA antisense strand are among the strategies used by *Squigualiser* to create the most intuitive visual representation of complex signal data. Currently, *Squigualiser* is a set of command line tools used to pre-process nanopore datasets and generate an interactive HTML browser snapshot, encompassing a specified region and set of aligned signal reads. Our future plans include development of a *Squigualiser* web server with a graphical interface to improve the user experience.

Above we outline two brief examples to illustrate how *Squigualiser* can be used to visually inspect, and thereby validate or interpret, candidate DNA/RNA modifications in ONT data. These highlight just one of many possible use cases for nanopore signal data visualisation. By providing the community with an efficient, intuitive visualisation framework, we anticipate *Squigualiser* will catalyse many new innovations and improvements in ONT signal data analysis. *Squigualiser* is freely available under an MIT licence: https://github.com/hiruna72/squigualiser

## METHODS & IMPLEMENTATION

### Squigualiser Usage

#### Installation

*Squigualiser* is developed and tested using Python 3.8 and can be installed via the pip package management tool. Prebuilt binaries (for linux-x86-64 and macOS-arm64 architectures), which can be simply extracted and executed, are released with each version. These pre-built binary packages were constructed using the snake-charming technique described in^24^, and contain the Python interpreter along with all the dependencies.

### Browser view

Squigualiser generates plots as HTML (or SVG) files that can be visualised on a modern web browser, such as Google Chrome or Microsoft Edge. The HTML plots have a toolbox that contains wheel/box zoom in/out, pan, freehand draw, save, and reset features that allow the user to interactively navigate the data and customise the display. The user can hover over a signal point to reveal its index value and the signal value. We provide a brief video guide to assist new users in navigating the *Squigualiser* browser view: https://youtu.be/kClYH4KpOjk. Squigualiser plots conform to several plot conventions to maintain reproducibility and consistency (**Supplementary Note 6**). Similar to the IGV, users can load an annotation file in BED format to assist in navigating the browser view. *Squigualiser* supports the more common 3-column BED format as well as the extended 12-column format. Hence, the user can visualise annotations in different colours, different tracks and with (or without) feature labels (**Supplementary Note 7**).

### Squigualiser reform and realign subtools

The subtools *reform* and *realign* are used for pre-processing the basecaller’s move table to be compatible with plotting subtools in *Squigualiser* (**Fig4a**). *Reform* takes an unaligned SAM/BAM file containing the move table and generates signal-to-read alignment in PAF format (containing *ss* tag). *Realign* takes the PAF output from *reform* along with read alignments in SAM format (e.g., Minimap2 alignments to the reference containing CIGAR string) to output the signal-to-reference alignment in SAM format. The following example commands demonstrate how the basecaller’s move table is pre-processed:

**Fig4.**
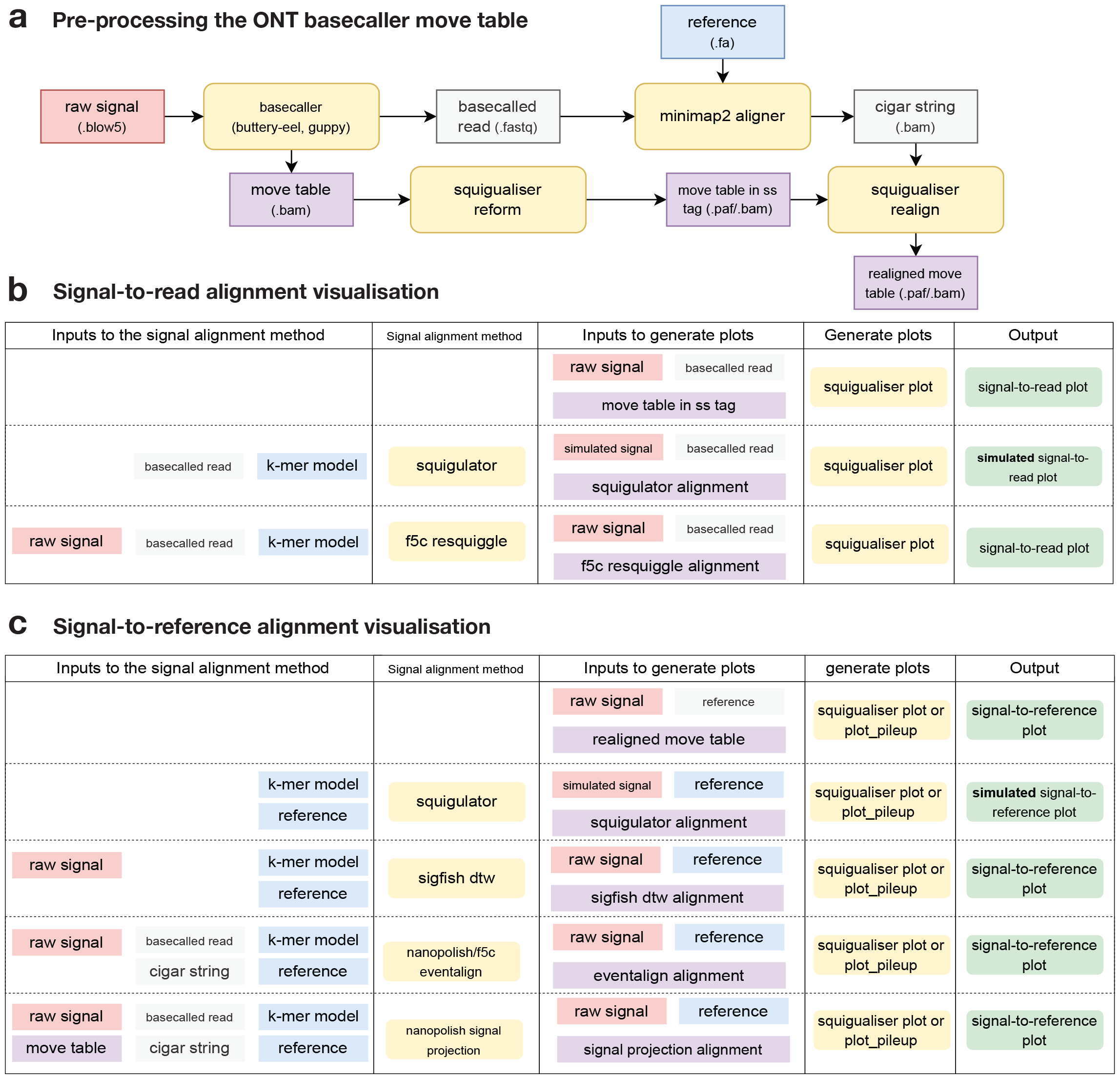
Overview of *Squigualiser* visualisation framework. Schematic diagram summarises data preprocessing steps and alternative workflow paths to signal alignment visualisation with *Squigualiser*. (**a**) The *calculate_offset* and *reform* subtools within *Squigualiser* may be used to convert the move table generated by ONT basecallers (*Guppy/Dorado*) into signal-to-read alignment format with *ss tag*s, which can be visualised directly, or can be translated to signal-to-reference alignments using the *realign* subtool (**b**). (**c**) Alternatively, the user may perform signal-to-reference alignment with a variety of external software, including *F5c, Nanopolish* or *Sigfish. Squigualiser plot* (single read) or *plot_pileup* (multiple reads in pileup format) are used to generate an interactive browser view (HTML format) capturing a specified set of reads and/or reference region, which can be displayed and navigated in a standard web browser (an example is provided in **Supplementary_File_1.html**). Our recent tool for signal data simulation, *Squigulator*, may be used to generate simulated signal-to-read and signal-to-reference alignments, which may be plotted as ‘expected data’ tracks by *Squigualiser*.

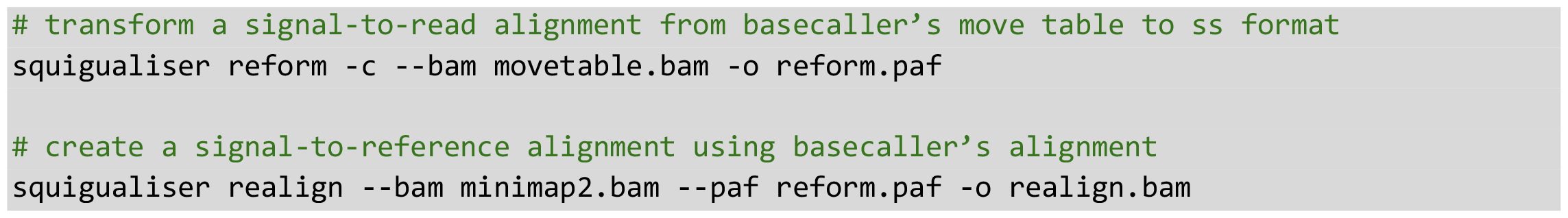

Plotting alignments from *f5c, Squigulator, Nanopolish* signal projection or *Sigfish* does not require any preprocessing, as we have modified these software to directly output in SAM format containing *ss* tag. The command-line options along with example commands for these tools to generate such output are described in the *Squigualiser* readme at https://hiruna72.github.io/squigualiser/.

### Squigualiser calculate_offsets subtool

The *calculate_offsets* subtool can be used to perform k-mer-to-base shift correction for more intuitive visualisation (discussed in detail in **Supplementary Note 3**). This subtool can be invoked either on a PAF file containing the *ss* tag (e.g., output from *reform*) or a k-mer model (e.g., *Nanopolish/f5c* k-mer model). k-mer-to-base shift correction can be performed using the following example command:

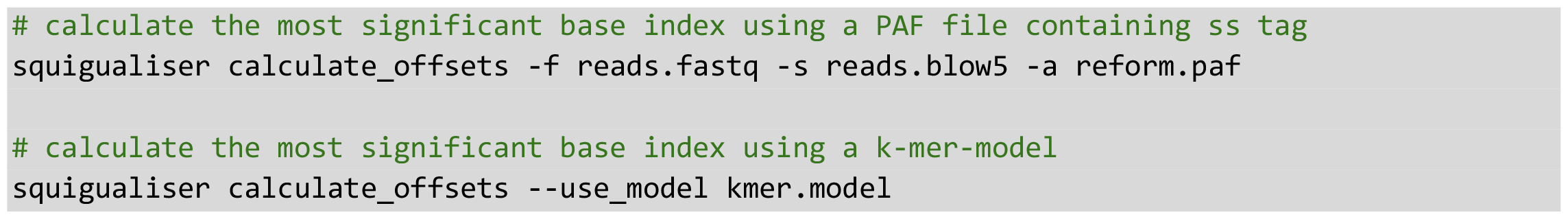

For the user’s convenience, the k-mer-to-base shift correction values for known k-mer models and basecaller models are provided as profiles (--*profile*) in *Squigualiser plot* and *plot_pileup* subtools (see **Supplementary Note 2, Table 2** and **Supplementary Note 3, Table 2**).

### Squigualiser plot, plot_pileup and plot_tracks subtools

Squigualiser has the subtools *plot, plot_pileup* and *plot_tracks* for plotting the data. The *plot* subtool is used to generate a plot for a single signal-to-read alignment or a single signal-to-reference alignment. For a signal-to-read plot, the inputs must be a PAF file containing ss tags (*reform, f5c resquiggle* or *Squigulator* PAF output), the raw signal file in BLOW5 format^25^ and the basecalled read in FASTQ format (**Fig4b**). For a signal-to-reference plot, the required inputs are a BLOW5 file, reference genome/transcriptome in FASTA format and signal-to-reference alignment in BAM/PAF format containing ss tags (**Fig4c**). This BAM/PAF file can be the output of *realign, f5c eventalign*, etc (**Fig4c**).

The *plot_pileup* subtool is used to generate signal-to-reference alignment plots in ‘pileup’ format, where individual reads are stacked vertically. The input for *plot_pileup* is the same as for the *plot* subtool above for signal-to-reference alignments (**Fig4c**). To visualise multiple pileups in a single HTML file (e.g., pileups from different samples or alternative alignment methods), the *plot_tracks* subtool can be used. *plot_tracks* accepts a text file with each *plot_pileup* command in a separate line.

The basic structure of the plotting commands is listed below.

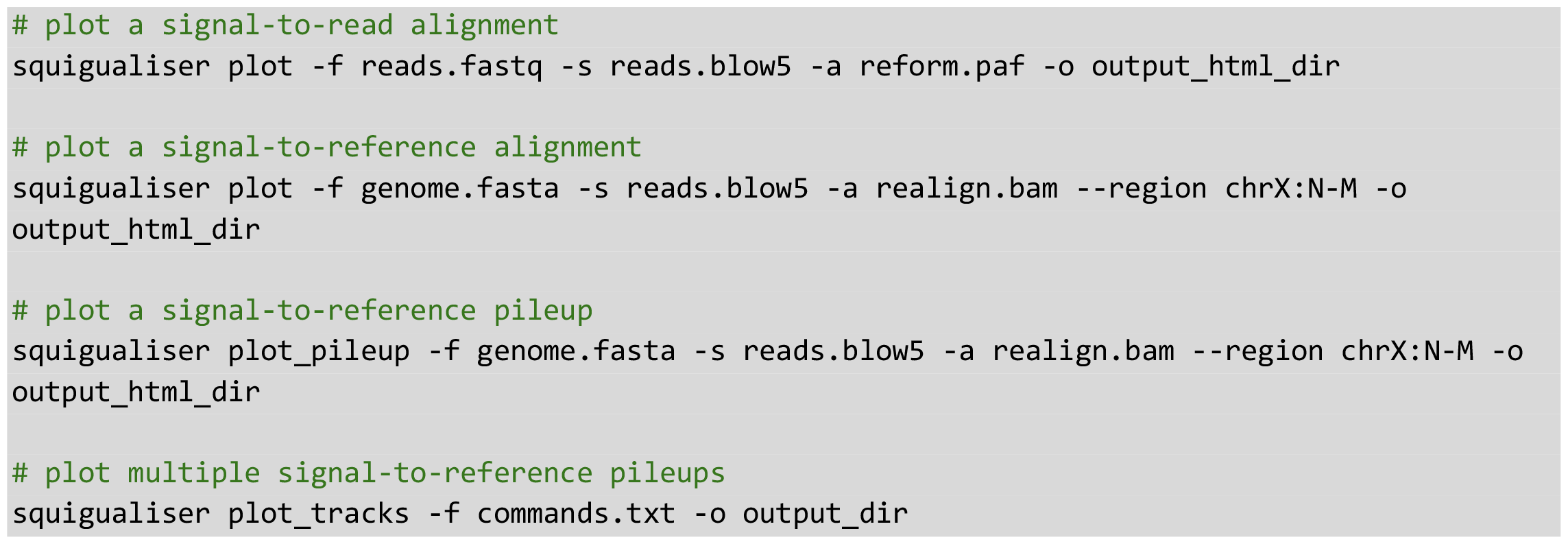

The user is provided with multiple data plotting and signal adjustment options in *plot, plot_pileup* and *plot_tracks* subtools. For example, the user can change the plot dimensions, initial x-range, hide base colours/signal samples etc. Further, the user can opt for pA conversion and several signal normalisation methods including Z-score scaling, Median-Median-Absolute-Difference scaling (med-MAD)^26^, and scaling to the k-mer model. In signal pileup mode the user can limit the number of signals to plot as well as the maximum number of signal points per signal. This will enhance user experience as the responsiveness of the plots can vary depending on the compute resources (GPU, Display resolution and size).

### Architecture of Squigualiser & Implementation

*Squigualiser* is developed in Python programming language and relies on several essential dependencies: *Bokeh* library to create interactive base-signal alignment plots; pyslow, r*eadpaf, pyfaidx, pyfastx*, and *pysam* for reading input file formats; *numpy* for data handling; and, *matplotlib* and *seaborn* to generate density plots in PDF format.

### Signal alignment String (SS) tag and output formats

Both a prerequisite and a motivation for developing *Squigualiser*, was to develop a standardised, optimised format for sequence-aligned nanopore signal data. Our signal alignment format is described in detail in **Supplementary Note 1**. Briefly, all the alignment operations (matches, deletions and insertions) can be encoded in a compact fashion using the *ss* tag. For example, ss:Z:7,2D3,4I,5 should be interpreted as 7 signal samples match, 2 bases deletion, 3 signal samples match, 4 signal samples insertion, followed by a 5 signal sample match along the sequence. This *ss* tag should be present as an auxiliary tag in PAF or SAM inputs to *Squigualiser*. For compatibility with *Squigualiser*, the PAF input columns 3, 4, 8 and 9 must correspond to the coordinates of *signal start, signal end, kmer start* and *kmer end*, respectively (**Supplementary Note 1**). In SAM format this information must be stored as a string with four comma-separated values in an auxiliary tag named *si*. The distinction between DNA and RNA signal alignments is made by swapping the values of *kmer start* and *kmer end* for RNA (**Supplementary Note 1**). Furthermore, two optional axillary tags named *sc* and *sh* may store the k-mer model calibration parameters to be used for scaling the signal to the k-mer model.

We modified the software packages *f5c, Sigfish, Squigulator and Nanopolish project-signal* subtool, to adhere to the aforementioned formats, thereby enabling their signal alignment outputs to be visualised in *Squigualiser*.

For basecallers from ONT that cannot be modified, we provide *Squigualiser* subtools *reform* and *realign* to convert the move table to *ss tag* format and translate the signal-to-read alignment to a signal-to-reference alignment respectively (see **Fig4a**). For example *reform* converts the move table mv:B:c:5:1,1,0,1,0,0,0,0,1,0,1,0,1,0 to the *ss* format as ss:Z:5,10,25,10,10,10. *Squigualiser realign* iterates through both the signal-to-read alignment generated by *reform* and the read-to-reference alignment, and edits the *ss* tag according to the CIGAR operations. Details on the move table format and how kmer-to-base shift correction is applied when generating the *ss* tag can be found in **Supplementary Notes 2 & 3**.

### k-mer-to-base shift correction

Many signal alignment methods that use k-mer models do not provide a signal-to-base alignment, instead provide a signal-to-k-mer alignment. To find the correct signal-to-base alignment we developed an algorithm below which is described in detail in **Supplementary Note 3**. Briefly, the algorithm applies a many-to-one mapping to create a 1-mer model that best reflects the properties of the particular k-mer model.

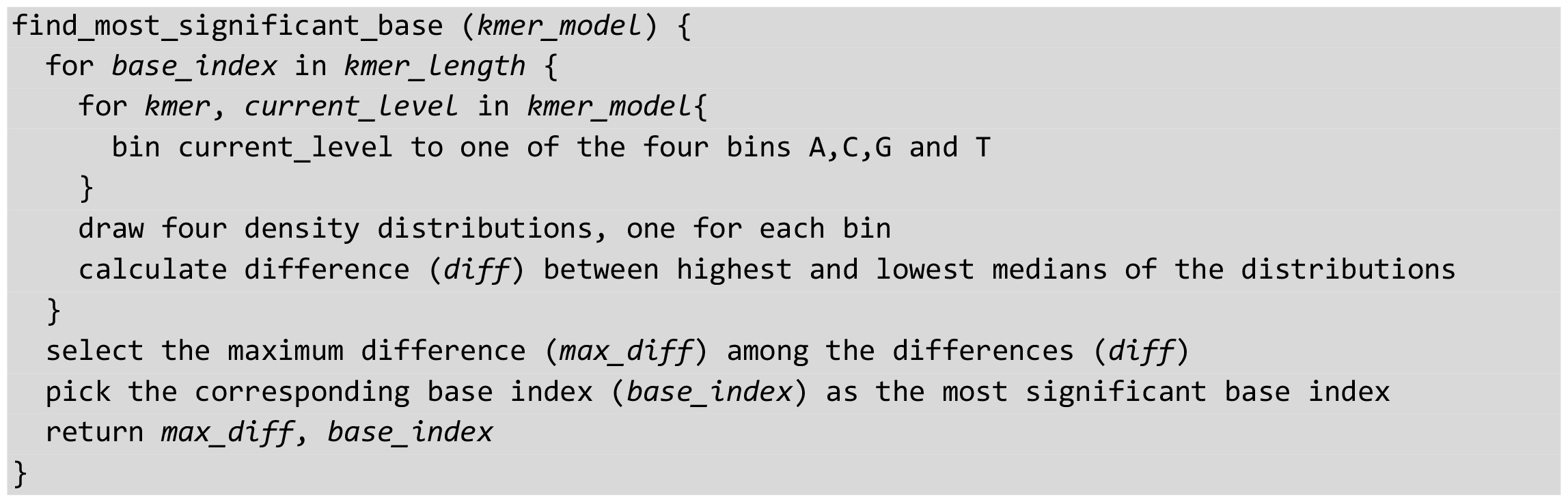

Even though the move table is a signal-to-base alignment, for some basecalling models it is not the optimum signal-to-base alignment. We treat the move table as an incorrect signal-to-1-mer alignment and adopt a correction method as above.

### Strategies behind fast region retrieval

If the user wants to plot a specific reference region, then the sequence file (in FASTA format) must be indexed (*faidx*) and the signal-to-sequence alignment (in SAM/PAF formats) file must be sorted and indexed. This is important as it is necessary to fetch only the necessary reads spanning across a given reference region. SAM format is preferred as it is well supported through *samtools* and its binary format (BAM) enables efficient region-based access. If PAF is used, the user has to sort it manually by providing the correct column indices for DNA/RNA, convert it to *bgzip* and index using *tabix*.

The raw signal which is the largest portion of the input data must be in BLOW5 format indexed using *slow5tools* ^27^, to enable efficient built-in random access functionality. The possibility to fetch only the necessary raw signals (random access) is vital to plot signal pileups spanning across a given reference region. Python package *pyslow5* is used to handle BLOW5 format files. These input formats enable *Squigualiser* to seamlessly generate signal pileup plots for a given reference region regardless of the size of the dataset.

The retrieval of aligned records for the user specified region (*ref_name:ref_start-ref_end*) from SAM/PAF file, followed by fetching the auxiliary tags (e.g., *ss* tag) for each alignment record, and then fetching the relevant raw signal from the SLOW5 file can be done as follows.

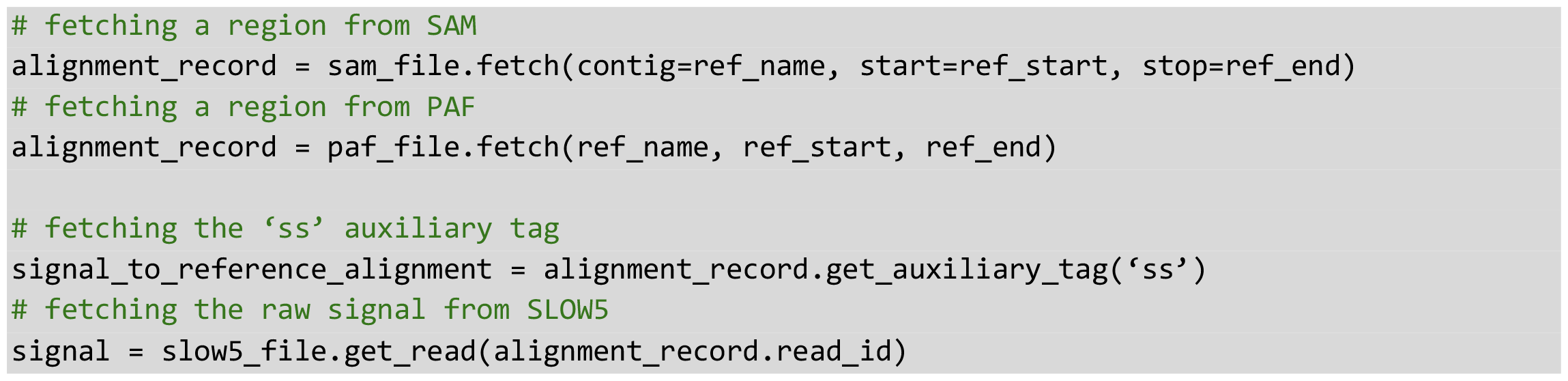

The fetched alignment_record is not guaranteed to exactly start and finish at the user-specified region. Instead, it can start before the user-specified start position and end before the user-specified end position. Hence, *Squigualiser* further refines the *alignment_record* and the *signal_to_reference_alignment* to get only the exact interval before generating plots.

## Supporting information

Supplementary_File_1.html

## ACKNOWLEDGEMENTS

We thank Jared Simpson for helping with the nanopolish k-mer pore model training and understanding *Nanopolish* signal projection; Sam Kovaka for developing and sharing *Uncalled4*, which inspired us to use interactive HTML plots in *Squigualiser*; and all users who have so far tested *Squigualiser* provided feedback or reported issues on GitHub.

We acknowledge the following funding support: Australian Medical Research Futures Fund grants MRF1173594, and MRF2023126 (to I.W.D.), Australian Research Council DECRA Fellowship DE230100178 (to H.G.) and Australian Research Council’s Discovery Project DP230100651 (to H.G and S.P). K.L. is supported by the Australian Government Research Training Program (RTP) Scholarship. The views expressed herein are those of the authors and are not necessarily those of the Australian Government or the Australian Research Council.

## AUTHOR CONTRIBUTIONS

H.S., H.G. and I.W.D. conceived and designed *Squigualiser*, devised the experiments and prepared the manuscript, with support from all authors. H.S. implemented Squigualiser, modified *Nanopolish* and conducted the experiments. H.G. designed the *ss* tag format and implemented the support for *F5c, Squigulator* and *Sigfish*. H.S. and I.W.D. generated the figures. All authors contributed to testing the software, read and approved the final manuscript.

## SUPPLEMENTARY MATERIAL

Supplementary Note 1: Signal Alignment Formats/Tags

Supplementary Note 2: Projecting basecaller move table to Signal alignment String (ss) tag Supplementary Note 3: Strategies behind signal alignment visualisation enhancements

Supplementary Note 4: Visualising a genomic region that has a CpG methylated site

Supplementary Note 5: Visualising RNA modifications

Supplementary Note 6: Plot conventions used by *Squigualiser*

Supplementary Note 7: Visualise BED annotations in *Squigualiser* plots

Supplementary_File_1.html: The signal-to-reference signal pileup plot is described in **Supplementary Note 4**.

## DECLARATIONS

I.W.D. manages a fee-for-service sequencing facility at the Garvan Institute of Medical Research and is a customer of Oxford Nanopore Technologies but has no further financial relationship. H.G., J.M.F. and I.W.D. have previously received travel and accommodation expenses from Oxford Nanopore Technologies. The authors declare no other competing financial or non-financial interests.

## DATA & CODE AVAILABILITY

The HG002 dataset sequenced on an ONT R10.4.1 PromethION flowcell used for the figures is publicly available on the NCBI Sequence Read Archive (SRR23215366). The synthetic unmodified and modified RNA datasets used in **Supplementary Note 5**, are also publicly available on NCBI SRA (SRR22888949, SRR22888950).

Squigualiser and all modified software associated with the manuscript is free and open source under an MIT licence:

*Squigualiser*: https://github.com/hiruna72/squigualiser

*F5c* with *ss tag* support: https://github.com/hasindu2008/f5c/releases/tag/v1.4

*Squigulator* with ss tag support: https://github.com/hasindu2008/squigulator/releases/tag/v0.2.2

*Sigfish* with ss tag support: https://github.com/hasindu2008/sigfish/releases/tag/v0.2.0-alpha

*Nanopolish* signal projection with ss tag support: https://github.com/hiruna72/nanopolish/tree/215592e0913418c2b20a86a67483cc1c6f557ca8

All the pipeline scripts, curated datasets, and generated plots can be found in the *Squigualiser* repository: https://github.com/hiruna72/squigualiser/tree/main/test/data/raw/pipelines

The user should be able to reproduce all our results by amending the scripts as explained in the associated documentation: https://hiruna72.github.io/squigualiser/docs/pipeline_basic

