## Supplementary Notes for "Interactive visualisation of raw nanopore signal data with Squigualiser"

### Supplementary Note 1: Signal Alignment Formats/Tags

Hiruna Samarakoon, Kisaru Liyanage, James M. Ferguson, Sri Parameswaran,
Hasindu Gamaarachchi, Ira W. Deveson

February 15, 2024

**Note: Read this document sequentially. Skipping sections without prior context is not recommended, as the content is not reiterated.**

This document explains the formats and tags supported by squigualiser. Signal-to-read alignment feature in Squigualiser requires the signal alignment information to be in PAF format explained in section 1.1. The signal-to-reference alignment feature can take either the PAF format (section 2.1) or SAM/BAM format (section 2.2). In both PAF and SAM format, the signal alignment is encoded using an auxiliary tag called the *Signal alignment String (ss)*. In PAF format, the start and end coordinates of the signal mapping are in relevant PAF columns, whereas in SAM format these coordinates are in the *Signal Information (si)* tag. Currently, the latest available versions of f5c and squigulator can directly output these formats compatible with squigualiser. We encourage developers working on signal alignment methods to adhere to this specification when generating outputs, ensuring wider compatibility with tools such as squigualiser.

### 1 Signal-to-read alignment

#### 1.1 PAF format

PAF format supported by Squigualiser (this format is inspired by UNCALLED) contains the columns described in Table 1. In this signal-to-read alignment context, the query is the raw-signal and the target is the basecalled-read.

Table 1: Columns in PAF format for signal-to-read alignment

| Col | Type | Name | Description |
|---|---|---|---|
| 1 | string | read_id | Read identifier name |
| 2 | int | len_raw_signal | Raw signal length (number of samples) |
| 3 | int | start_raw | Raw signal start index (0-based; BED-like; closed) |
| 4 | int | end_raw | Raw signal end index (0-based; BED-like; open) |
| 5 | char | strand | Relative strand: "+" or "-" |
| 6 | string | read_id | Same as column 1 |
| 7 | int | len_kmer | base-called sequence length (no. of k-mers) |
| 8 | int | start_kmer | k-mer start index on basecalled sequence (0-based; see note below) |
| 9 | int | end_kmer | k-mer end index on basecalled sequence (0-based; see note below) |
| 10 | int | matches | Number of k-mers matched on basecalled sequence |
| 11 | int | len_block | Same as column 7 |
| 12 | int | mapq | Mapping quality (0-255; 255 for missing) |

**Conventions:** For DNA reads, column 8 is a closed coordinate (inclusive) and column 9 is an open coordinate (not-inclusive); and, column 8 coordinate is always smaller than column 9. However, this is different in direct-RNA reads because the sequencing of direct-RNA happens in the reverse direction (3'->5') and therefore the raw signal is also in the reverse direction. The basecalled read output by basecallers is however in the correct direction (5'->3'). Thus, For RNA reads, the column 8 coordinate will be larger than that of column 9. Column 9 is a closed coordinate (inclusive) while column 8 is an open coordinate (not-inclusive) in this case, contrary to DNA. Columns 10,11 and 12 are not used by Squigualiser and thus are not finalised.

**Auxiliary tags:** Auxiliary tags supported by squigualiser are described in Table 2, where the *ss* tag is mandatory. The `sh` and `sc` tag values can be used to scale the raw signal to the pore model. This can be done as: `scaled pA current values = (pA - sh) / sc`, where, `pA = (raw_signal + offset) * range / digitisation)`. The *ss* tag is described in detail below in section 1.1.1

Table 2: Auxilliary tags in PAF format

| Tag | Type | Description |
|---|---|---|
| sc | f | Post alignment recalibrated scale parameter |
| sh | f | Post alignment recalibrated shift parameter |
| ss | Z | signal alignment string in format |

##### 1.1.1 ss tag

*ss* tag is a custom encoding that compacts the signal-base alignment. It can be thought of as an extended version CIGAR string that accommodates for signal alignment needs. This *ss* string was inspired by the *--sam* option in Nanopolish for eventalign.

Consider the example `8,5,4,8I4,3D4,5,` for DNA. This means 8 signal samples map to the starting base of the sequence; the next 5 samples to the next base, the next 4 samples to the next base; 8 next samples are missing a mapping in the basecalled read (insertion to reference); 4 samples map to the next base; 3 bases in the basecalled read have no corresponding signal samples (deletion); 4 samples map to the next base; and 5 samples map to the next base. Note that the start indexes of read and signal are the absolute values in columns 8 and 3 respectively above. the *ss* string is relative to this.

The '`,`', '`D`' and '`I`' can be though of as three different operations. The '`,`' after a number means step one base in the basecall reference, while step number of samples preceding the '`,`' in the raw signal. The '`D`' after a number means step number of bases preceding the '`D`' in the basecalled read and no stepping in the raw signal. The '`I`' after a number means step number of samples preceding the '`I`' in the raw signal and no stepping in the basecalled read.

##### 1.1.2 DNA Example

To make things further clear, given in Figure 1 is an illustration for an alignment of DNA raw-signal to a basecalled read.

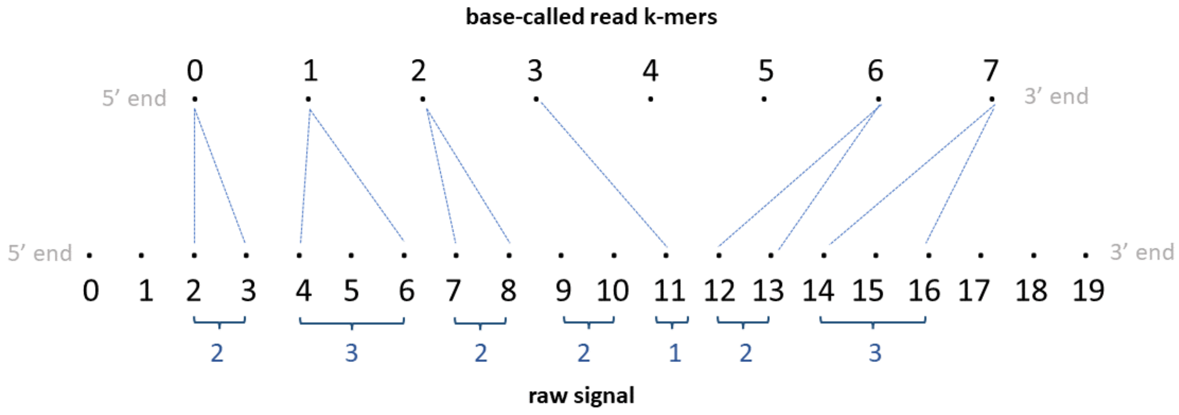


Figure 1: DNA example for signal-to-read alignment

Each dot in the top sequence in Figure 1, represents a k-mer and the number above each dot is the corresponding k-mer index. If the basecalled read is `ACGGTAACTATAC` and assuming the k-mer size in the k-mer model is 6, the 0th k-mer is `ACGGTA`, 1st k-mer is `CGGTAA` ... and the 7th k-mer is `CTATAC`. Each dot in the bottom sequence in the illustration represent a raw-signal sample and the number below each dot is the corresponding signal index.

The PAF output for the example in Figure 1 will look like in Table 3 (the header is not present in the actual output).

Table 3: DNA signal-to-read alignment example in PAF format

| read_id | len_raw_signal | start_raw | end_raw | strand | read_id | len_kmer | start_kmer | end_kmer | matches | len_block | mapq | |
|---|---|---|---|---|---|---|---|---|---|---|---|---|
| rid0 | 20 | 2 | 17 | + | rid0 | 8 | 0 | 8 | 6 | 8 | 255 | ss:Z:2,3,2,2I1,2D2,3, |

##### 1.1.3 direct-RNA Example

Now see the illustration in Figure 2 for direct-RNA.

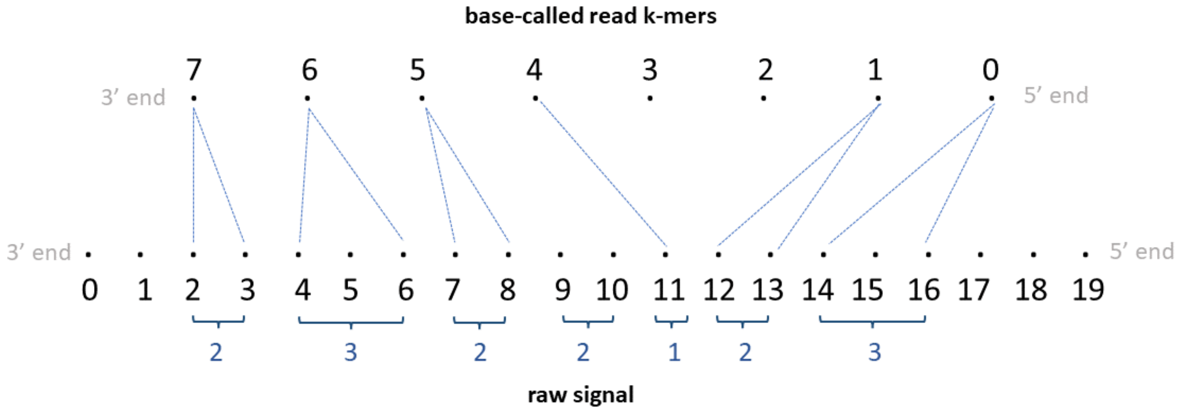


Figure 2: RNA example for signal-to-read alignment

Note that the RNA is sequenced from 3'->5' end, so the raw signal is 3'->5' direction. As the basecaller outputs the basecalled read in 5'->3' direction, the basecalled read is reversed to be 3'->5' in the illustration (note: indices in illustration denote the actual index in the basecalled read in 5'->3' direction). If the basecalled read in 5'->3' direction is `ACGGUAACUAUA` and assuming the k-mer size in the k-mer model is 5, the 0th k-mer is `ACGGU`, 1st k-mer is `CGGUA` . . . and the 7th k-mer is `CUAUA`.

The PAF output for the example RNA in Figure 2 will look like in Table 4.

Table 4: RNA signal-to-read alignment example in PAF format

| read_id | len_raw_signal | start_raw | end_raw | strand | read_id | len_kmer | start_kmer | end_kmer | matches | len_block | mapq | |
|---|---|---|---|---|---|---|---|---|---|---|---|---|
| rid0 | 20 | 2 | 17 | + | rid0 | 8 | 8 | 0 | 6 | 8 | 255 | ss:Z:2,3,2,2I1,2D2,3, |

##### 1.1.4 C code snippet to parse ss tag

A C code snippet that converts the value in the ss tag (a readable code that is not optimised) is given below:

```c
#include <stdio.h>
#include <stdlib.h>
#include <ctype.h>

#define MAX_LEN_KMER 20000

int main(){

    //paf fields
    char *ss="2,3,2,2I1,2D2,3,";
    int start_raw=2; int end_raw=17; int len_raw_signal=20;
    int start_kmer=0; int end_kmer=8; int len_kmer=8;

    // Raw signal start index for the corresponding k-mer and Raw signal end index for the corresponding k-mer
    int st_raw_idx[MAX_LEN_KMER]; int end_raw_idx[MAX_LEN_KMER];

    //intialise to -1
    for(int i=0; i<MAX_LEN_KMER; i++){ st_raw_idx[i]=end_raw_idx[i]=-1; }

    int st_k = start_kmer; int end_k = end_kmer; //if DNA, start k-kmer index is start_kmer column in paf and end k-kmer index is end_kmer column in paf
    int8_t rna = start_kmer > end_kmer ? 1 : 0; //if RNA start_kmer>end_kmer in paf
    if(rna){ st_k = end_kmer; end_k = start_kmer; } //if RNA, start k-kmer index is end_kmer column in paf and end k-kmer index is start_kmer column in paf

    int i_k = st_k; int i_raw = start_raw; //current k-mer index and current raw signal index

    //buffer for storing digits preceding each operation and its index
    char buff[11]; int i_buff=0;

    while(*ss){

        if(*ss==',' || *ss=='I' || *ss=='D'){
            if(i_buff <= 0){ fprintf(stderr,"Bad ss: Preceding digit missing\n"); exit(1); }//if nothing in buff

            buff[i_buff]=0; //null terminate buff
            int num = atoi(buff);
            if(num < 0){ fprintf(stderr,"Bad ss: Cannot have negative numbers\n"); exit(1); }
            i_buff=0; buff[0]=0; //reset buff

            if(*ss=='I'){ //if an insertion, current raw signal index is incremented by num
                i_raw += num;
            } else if(*ss=='D'){ //if an deletion, current k-mer index is incremented by num
                i_k += num;
            } else if (*ss==','){ //if a mapping, increment accordingly and set raw signal indices for the current k-mer
                end_raw_idx[i_k] = i_raw; i_raw += num;
                st_raw_idx[i_k] = i_raw; i_k++;
            }
        } else {
            if(!isdigit(*ss)){ fprintf(stderr,"Bad ss: A non-digit found when expected a digit\n"); exit(1); }
            buff[i_buff++]=*ss;
        }
        ss++;
    }

    if(i_raw!=end_raw){ fprintf(stderr,"Bad ss: Signal end mismatch\n"); exit(1); } // current raw signal index should be equal to end_raw
    if(i_k!=end_k){ fprintf(stderr,"Bad ss: Kmer end mismatch\n"); exit(1); } //current k-mer index should be equal to end_k

    for(int i=st_k; i<end_k; i++){
        if(end_raw_idx[i]==-1){
            if(st_raw_idx[i] != -1) { fprintf(stderr,"Bad ss: This shoud not have happened\n"); exit(1); }//if st_raw_idx[i] is -1, then end_raw_idx[i] should also be -1
```

```
            printf("%d\t.\t.\n", rna ? len_kmer-i-1 : i);
        }else {
            printf("%d\t%d\t%d\n", rna ? len_kmer-i-1 : i, end_raw_idx[i], st_raw_idx[i])
    ;
        }
    }
}
```

### 2 Signal-to-reference alignment

#### 2.1 PAF format

The PAF format supported by Squigualiser for the signal-to-reference feature is similar to PAF output explained in section 1.1, with major difference being that the "basecalled read" is now the "refererence sequence". Assuming that the reader is well familiarised with the PAF output explained in section 1.1, that information is not repeated here, instead only a summary is given. The PAF columns are explained in Table 5. Unlike in section 1.1, the strand column (column 5) can be now both '+' and '-'. The query is the raw-signal and the target is the reference.

Table 5: Columns in PAF format for signal-to-reference alignment

| Col | Type | Name | Description |
|---|---|---|---|
| 1 | string | read_id | Read identifier name |
| 2 | int | len_raw_signal | Raw signal length (number of samples) |
| 3 | int | start_raw | Raw signal start index (0-based; BED-like; closed) |
| 4 | int | end_raw | Raw signal end index (0-based; BED-like; open) |
| 5 | char | strand | Relative strand: "+" or "-" |
| 6 | string | read_id | Reference sequence name |
| 7 | int | len_kmer | Reference sequence length (no. of k-mers) |
| 8 | int | start_kmer | k-mer start index on reference sequence (0-based; see note below) |
| 9 | int | end_kmer | k-mer end index on sequence sequence (0-based; see note below) |
| 10 | int | matches | Number of k-mers matched on reference sequence |
| 11 | int | len_block | Number of k-mers on the mapped segment on reference sequence |
| 12 | int | mapq | Mapping quality (0-255; 255 for missing) |

The conventions (paragraph conventions) and auxiliary tags (paragraph auxiliary tags and section 1.1.1 described before are applicable here. Some examples are given below.

##### 2.1.1 DNA examples

**DNA Positive strand example:** Assume we have a read signal named rid0 of 1000 signal samples, mapped to a reference contig named ctg0 of 35 bases Assume a k-mer size of 6. We have a total of 30 k-mers in the reference. Assume the signal-reference alignment looks like in Figure 3. Assume that the 12-24th bases (0-index; bed-like) in this contig are `TTGATGGTGGAA`. Thus, 12th kmer is TTGATG, 13th k-mer is TGATGG, 14th k-mer is GATGGT, .. and the 18th k-mer is GTGGAA.

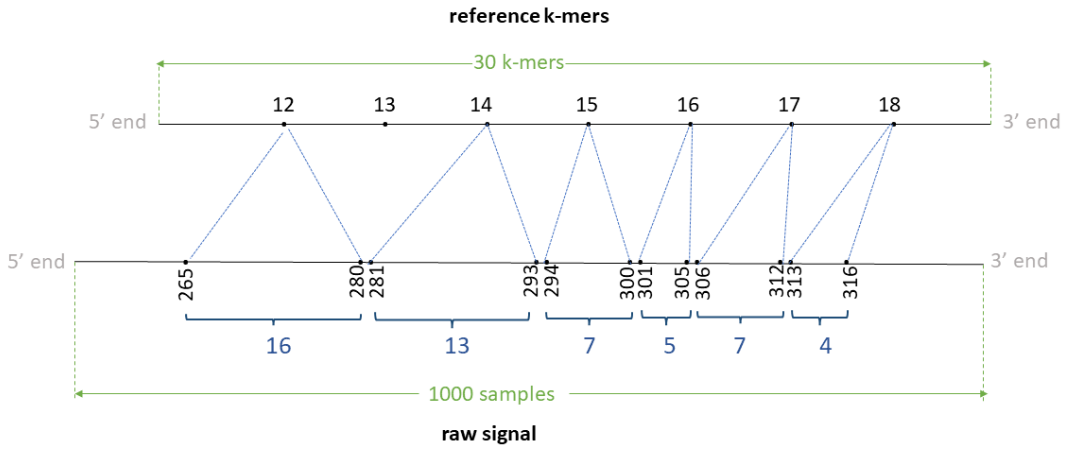


Figure 3: DNA example for signal-to-reference alignment (mapped to positive strand)

The PAF output from eventalign will look like in Table 6 (the header is not present in the actual output).

Table 6: DNA signal-to-reference alignment example in PAF format (mapped to positive strand)

| read_id | len_raw_signal | start_raw | end_raw | strand | ref_id | len_kmer | start_kmer | end_kmer | matches | len_block | mapq | |
|---|---|---|---|---|---|---|---|---|---|---|---|---|
| rid0 | 1000 | 265 | 317 | + | ctg0 | 30 | 12 | 19 | 6 | 7 | 255 | ss:Z:16,1D13,7,5,7,4, |

**DNA Negative strand example:** Assume we have a read signal named rid1 of 1000 signal samples, mapped to a reference contig named ctg0 of 35 bases Assume a k-mer size of 6. We have a total of 30 k-mers in the reference. Assume the signal-reference alignment looks like in Figure 4 (note: indices in illustration denote the actual index in the + strand of the reference genome in 5'->3' direction).

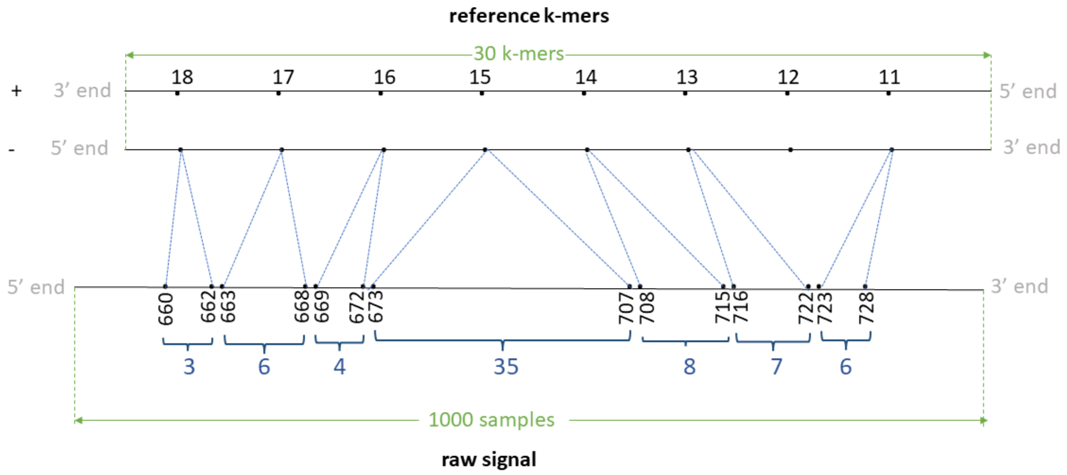


Figure 4: DNA example for signal-to-reference alignment (mapped to negative strand)

Assume that the 11-24th bases (0-index; bed-like) in this contig are ATTGATGGTGGAA. Thus, 11th kmer is ATTGAT, 12th k-mer is TTGATG, 13th k-mer is TGATGG, .. 17th k-mer is GGTGGA and the 18th k-mer is GTGGAA. Th negative strand is like:

```
5' ATTGATGGTGGAA 3' + strand
   |||||||||||||
3' TAACTACCACCTT 5' - strand
```

The Reverse complement is thus TTCCACCATCAAT. The 11th k-mer ATTGAT in the + strand relates to ATCAAT in the - strand, 12th k-mer TTGATG relates to CATCAA, 13th k-mer TGATGG relates to CCATC ,. . . , 17th k-mer GGTGGA relates to TCCACC and 18th k-mer GTGGAA relates to TTCCAC.

The PAF output from eventalign will look like in Table 7 (the header is not present in the actual output).

Table 7: DNA signal-to-reference alignment example in PAF format (mapped to negative strand)

| read_id | len_raw_signal | start_raw | end_raw | strand | ref_id | len_kmer | start_kmer | end_kmer | matches | len_block | mapq | |
|---|---|---|---|---|---|---|---|---|---|---|---|---|
| rid1 | 1000 | 660 | 729 | - | ctg0 | 30 | 11 | 19 | 7 | 8 | 255 | ss:Z:3,6,4,35,8,7,1D6, |

##### 2.1.2 RNA examples

**RNA Positive strand example:** Assume we have a read signal named rid0 of 3000 signal samples, mapped to a reference transcript (or can be a ctg in the reference genome) named trn0 of 65 bases. Assume a k-mer size of 5. We have a total of 61 k-mers in the reference.

Assume the signal-reference alignment looks like in the Figure 5. Note that the RNA is sequenced 3'->5' end, so the raw signal is 3'->5' direction. However, as transcripts in the reference are in 5'->3' direction, the transcript is reversed to be 3'->5' in the illustration (note: indices in illustration denote the actual index in the transcript in 5'->3' direction).

Assume that the 45-56th bases (0-index; bed-like) in this transcript in 5'->3' direction is GAGAGCCCTGA. Then, 45th kmer is GAGAG, 46th k-mer is AGAGC, 47th k-mer is GAGCC, .. and the 51st k-mer is CCTGA.

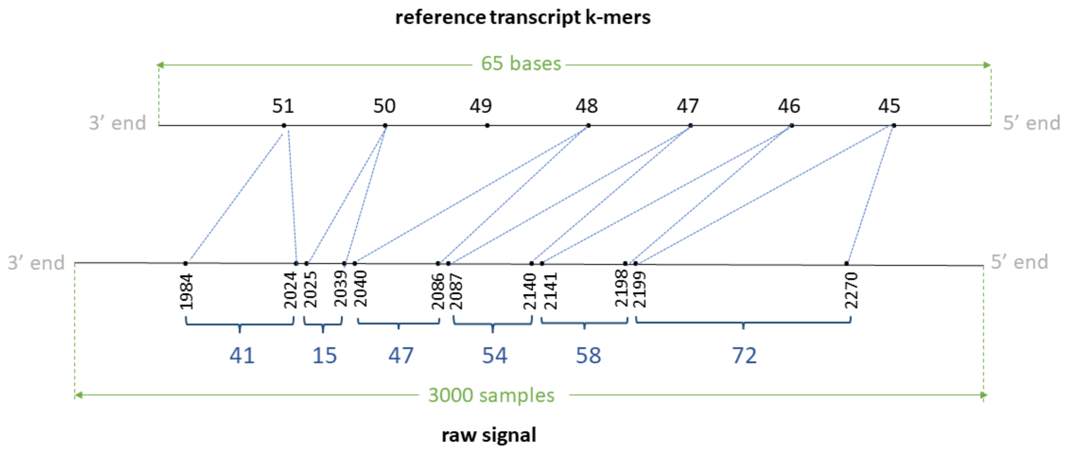


Figure 5: RNA example for signal-to-reference alignment (mapped to positive strand)

The PAF output from eventalign will look like in Table 8 (the header is not present in the actual output):

Table 8: RNA signal-to-reference alignment example in PAF format (mapped to positive strand)

| read_id | len_raw_signal | start_raw | end_raw | strand | ref_id | len_kmer | start_kmer | end_kmer | matches | len_block | mapq | |
|---|---|---|---|---|---|---|---|---|---|---|---|---|
| rid0 | 3000 | 1984 | 2271 | + | trn0 | 61 | 52 | 45 | 6 | 7 | 255 | ss:Z:41,15,1D47,54,58,72, |

Note that start_kmer and end_kmer are otherway round compared to DNA.

**RNA Negative strand example** Assume we have a read signal named rid1 of 500 signal samples, mapped to a reference contig named ctg1 of 20 bases. Assume a k-mer size of 5. We have a total of 24 k-mers in the reference. Assume the signal-reference alignment looks like in Figure 6 (note: indices in illustration denote the actual index in the + strand of the reference genome in 5'->3' direction).

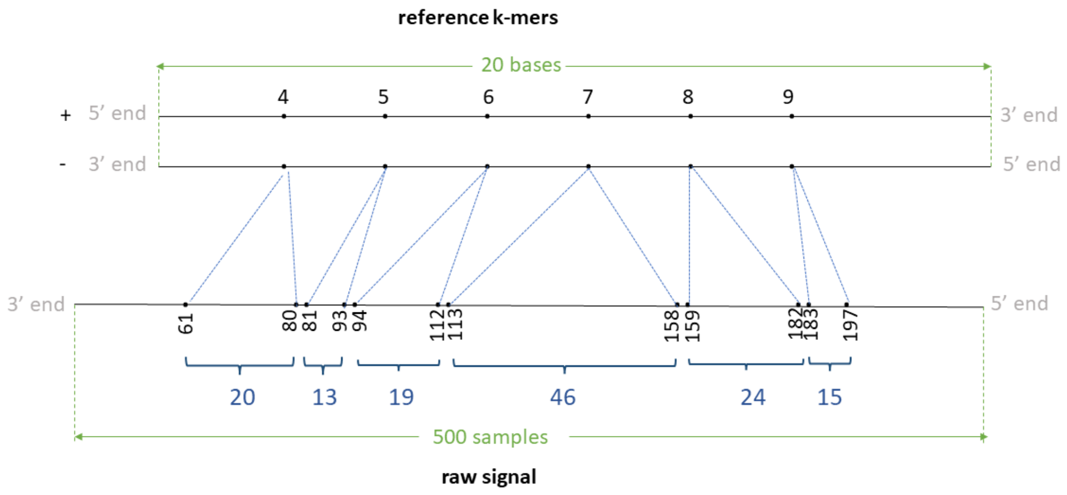


Figure 6: RNA example for signal-to-reference alignment (mapped to negative strand)

Assume that the 4-13th bases (0-index; bed-like) in this contig are AAATGGCTGA. Thus, 4th kmer is AAATG, 5th k-mer is AATGG, .. 8th k-mer is GGCTG and the 9th k-mer is GCTGA. The negative strand is like:

```
5' AAATGGCTGA 3' + strand
   ||||||||||
3' TTTACCGACT 5' - strand
```

The Reverse complement is thus TCAGCCATTT. The 4th k-mer AAATG in the + strand relates to CATTT in the - strand, 5th k-mer AATGG relates to CCATT ,... , 8th k-mer GGCTG relates to CAGCC and 9th k-mer GCTGA relates to TCAGC.

The PAF output from eventalign will look like in Table 9 (the header is not present in the actual output):

Table 9: RNA signal-to-reference alignment example in PAF format (mapped to negative strand)

| read_id | len_raw_signal | start_raw | end_raw | strand | ref_id | len_kmer | start_kmer | end_kmer | matches | len_block | mapq | |
|---|---|---|---|---|---|---|---|---|---|---|---|---|
| rid1 | 500 | 61 | 198 | - | ctg1 | 24 | 10 | 4 | 6 | 6 | 255 | ss:Z:20,13,19,46,24,15, |

#### 2.2 SAM format

Auxiliary tags supported by squigualiser in SAM format are described in Table 10, where the *si* and *ss* tags are mandatory. The `sh` and `sc` tag values can be used to scale the raw signal to the pore model in the same way stated in paragraph auxiliary tags for PAF format. The *si* tag contains four comma separated values *start_raw*, *end_raw*, *start_kmer* and *end_kmer*, respectively. Those values are the same as the columns 3,4,8 and 9 in the PAF format explained in Table 5 for PAF format. The *ss* tag is same as described in detail in section 1.1.1.

Table 10: Auxilliary tags in SAM format

| Tag | Type | Description |
|---|---|---|
| sc | f | Post alignment recalibrated scale parameter |
| sh | f | Post alignment recalibrated shift parameter |
| si | Z | signal information tag containing coordinates associated with the ss tag |
| ss | Z | signal alignment string in format described under section 1.1.1 |

### Supplementary Note 2: Projecting basecaller move table to Signal alignment String (ss) tag

Hiruna Samarakoon, Kisaru Liyanage, James M. Ferguson, Sri Parameswaran, Hasindu Gamaarachchi, Ira W. Deveson

February 15, 2024

This document explains the format of the move table generated by the ONT basecallers (Section 1), how the move table is projected to our Signal alignment String (ss) format (Section 2), and how a preliminary signal-to-reference alignment is obtained using the move table information (Section 3). The move table is a signal-to-read alignment format introduced by Oxford Nanopore Technologies (ONT), lacking public documentation. We've compiled this documentation based on available information, recognising that ONT may change the format in the future. In order to support move table based alignments in *Squigualiser*, *reform* subtool converts move table to ss tag. SS tag is documented in **Supplementary Note 1**. *Squigualiser's realign* subtool can calculate a signal-to-reference alignment in conjunction with the read-to-reference alignment information.

#### 1 Move table

ONT basecallers output move arrays in SAM/BAM format. The important fields are listed below.

1. Primary field - `read_id`
2. Primary field - basecalled fastq sequence length
3. Primary field - basecalled fastq sequence
4. Auxiliary tag 'ns' - raw signal length
5. Auxiliary tag 'ts' - raw signal trim offset
6. Auxiliary tag 'mv' - move table

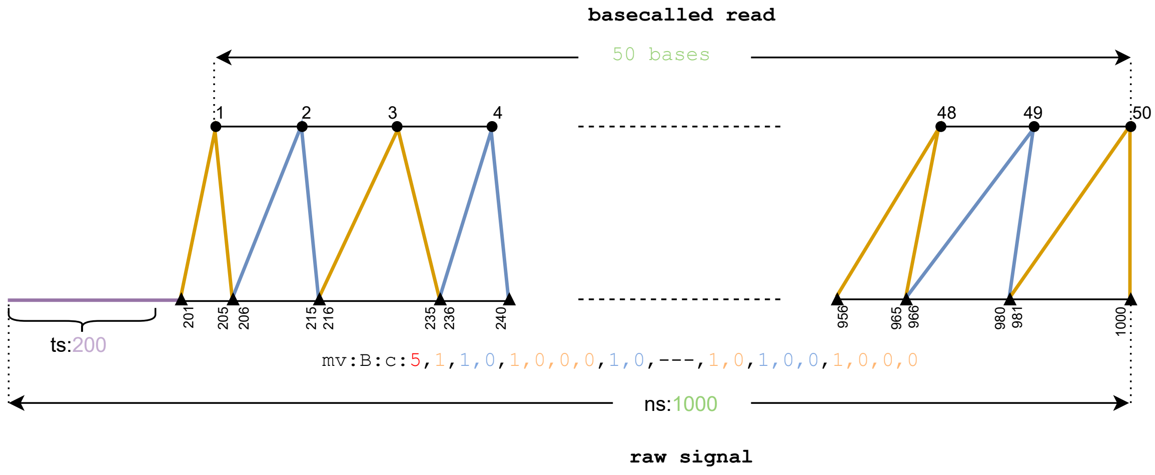


Figure 1: Move table example for signal-to-read alignment

An example move table looks like the following where 'mv' is the tag name and 'B:c:' denotes an array of 'int8_t' elements.

$$\text{mv:B:c:}5, 1, 1, 0, 1, 0, 0, 0, 1, 0, 1, 0, 1, 0, 0, 0, 1, 0, 1, 0, 1, 1, 0, 1, 0, 1, 0, 1, 1, 1, 1, \ldots$$

The downsampling factor used in the neural network (stride) is always the first integer. In the above case it is 5. The rest is the actual move array. The number of ones (1) in the move array equals the fastq sequence length. According to the above example the first move corresponds with 1 x stride signal points. The second move corresponds with 2 x stride signal points. The third with 4 x stride, the fourth with 2 x stride and so on (Figure 1).

The basecalling models have different stride values. The stride values extracted from the move tables generated by Guppy 6.5.7 are listed in Table 1.

Table 1: Strides values of some of the guppy basecaller (v6.5.7) models

| Model | Stride |
|---|---|
| dna_r10.4.1_e8.2_400bps_fast.cfg | 5 |
| dna_r10.4.1_e8.2_400bps_hac.cfg | 5 |
| dna_r10.4.1_e8.2_400bps_sup.cfg | 5 |
| dna_r9.4.1_450bps_fast_prom.cfg | 5 |
| dna_r9.4.1_450bps_hac_prom.cfg | 5 |
| dna_r9.4.1_450bps_sup_prom.cfg | 5 |
| rna_r9.4.1_70bps_hac_prom.cfg | 10 |
| rna_r9.4.1_70bps_fast_prom.cfg | 12 |

#### 2 *Squigualiser reform* subtool

The move table format is not general format. A move that is not a multiple of the stride and indels cannot be stored in the move table format. Moreover, it can only encode signal-to-read alignments and cannot encode signal-to-reference alignments. A generalised format called ss tag that we introduced is documented in **Supplementary Note 1**. For example, this move string mv:B:c:5:1,1,0,1,0,0,0,0,1,0,1,0,1,0 can be stored in the ss tag format as ss:Z:5,10,25,10,10,10. Furthermore, complex alignments can be stored in the ss tag format. For example, ss:Z:7,2D3,4I,5 should be interpreted as 7 samples match, 2 bases deletion, 3 samples match, 4 samples insertion, followed by a 5 samples match. Such alignments are not directly possible to be encoded in the current version of the move table format.

Therefore, input alignments should be in the ss tag format to generate plots using *Squigualiser*, i.e. move table must be converted to the ss tag format. This is done using *Squigualiser reform* subtool. The two important parameters *reform* takes are *kmer_length* (or *k*) and *sig_move_offset* (or *m*). This determines the best base colour adjustment to the signal events (**Supplementary Note 3**). The user can provide the *--profile* parameter to use predetermined values *kmer_length* and *sig_move_offset* (Table 2).

Table 2: Predetermined *kmer_length* and *sig_move_offset* values for the guppy basecaller (v6.5.7) models

| DNA/RNA | Profile Name | Kmer Length | Sig Move Offset |
|---|---|---|---|
| DNA | guppy_dna_r9.4.1_450bps_fast | 3 | 2 |
| DNA | guppy_dna_r9.4.1_450bps_fast_prom | 3 | 2 |
| DNA | guppy_dna_r9.4.1_450bps_hac | 3 | 2 |
| DNA | guppy_dna_r9.4.1_450bps_hac_prom | 3 | 2 |
| DNA | guppy_dna_r9.4.1_450bps_sup | 4 | 3 |
| DNA | guppy_dna_r9.4.1_450bps_sup_prom | 4 | 3 |
| DNA | guppy_dna_r10.4.1_e8.2_400bps_fast | 2 | 1 |
| DNA | guppy_dna_r10.4.1_e8.2_400bps_fast_prom | 2 | 1 |
| DNA | guppy_dna_r10.4.1_e8.2_400bps_hac | 2 | 1 |
| DNA | guppy_dna_r10.4.1_e8.2_400bps_hac_prom | 2 | 1 |
| DNA | guppy_dna_r10.4.1_e8.2_400bps_sup | 2 | 1 |
| RNA | guppy_rna_r9.4.1_70bps_fast | 1 | 0 |
| RNA | guppy_rna_r9.4.1_70bps_fast_prom | 1 | 0 |
| RNA | guppy_rna_r9.4.1_70bps_hac | 1 | 0 |
| RNA | guppy_rna_r9.4.1_70bps_hac_prom | 1 | 0 |

### 3 *Squigualiser realign* subtool

As discussed before, *Squigualiser reform* converts the basecaller move table to ss format. Subsequently, employing the read-to-reference alignment (CIGAR string) enables the derivation of a signal-to-reference alignment. This is implemented in *Squigualiser realign* subtool as explained in Figure 2 and in Algorithm 1. *Squigualiser realign* subtool produces the signal-to-reference alignment in ss tag format

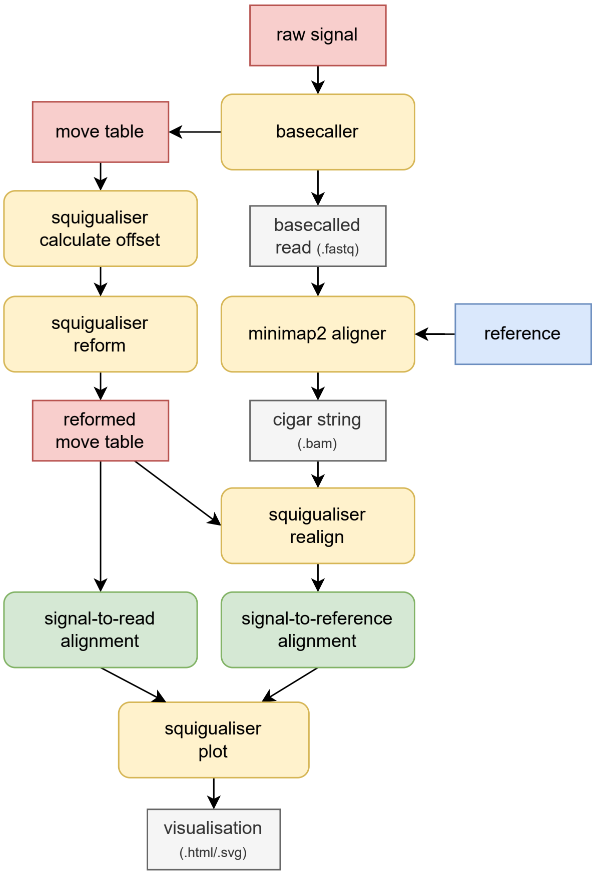


Figure 2: *Squigualiser reform* and *realign* subtools are used for signal-to-read alignment and signal-to-reference alignment respectively.

**Algorithm 1:** *Squigualiser realign* subtool

**Data:** CIGAR string (read-to-reference alignment), reformed array (signal-to-read alignment)
**Result:** reformed array (signal-to-reference alignment)

```
for each position in CIGAR string and reformed array do
    // Process soft-clipped bases
    if current position corresponds to soft-clipped bases then
        Clip corresponding moves from the reformed array;
    // Process insertions
    if current position corresponds to an insertion ('I' in CIGAR string) then
        Add an 'I' operator to the reformed array;
    // Process deletions
    if current position corresponds to a deletion ('D' in CIGAR string) then
        Add a 'D' operator to the reformed array;
    // Handle other CIGAR operations accordingly
    // Handle DNA/RNA accordingly
```

### Supplementary Note 3: Strategies behind signal alignment visualisation enhancements

Hiruna Samarakoon, Kisaru Liyanage, James M. Ferguson, Sri Parameswaran, Hasindu Gamaarachchi, Ira W. Deveson

February 19, 2024

**Note: Read this document sequentially. Skipping sections without prior context is not recommended, as the content is not reiterated.**

This document first explains the concept of k-mer model and the ***most significant base*** in Section 1. Then, Section 2 presents the technical details about calculating the most significant base. Section 3 demonstrates how the correct application of the most significant base concept can be further validated using an example involving heterozygous A/T SNV.

#### 1 K-mer model & most significant base

Different nanopore chemistries have different pore specifications, thus, different k-mer lengths, current levels and standard deviations. This information is stored in a file called k-mer model/table (also known as pore model/table). Many signal alignment methods including *Nanopolish/f5c eventalign, Nanopolish signal projection, Squigulator simulation*, and *Sigfish dtw* use k-mer models. At a given moment, all the bases of the k-mer inside the nanopore influence the current/voltage level. However, not all the bases equally contribute to this phenomenon (Fig. 1). Our aim is to visualise the most significant base (that mostly influences the current level) and the relevant signal event together.

Depending on the method used to generate the k-mer model, the central base of the k-mer may or may not be the most significant base (Table 1). Given a pore-model, we can find the position of the most significant base (that influences the k-mer current level the most) in the k-mer, by analysing the degree to which each base in the k-mer can discriminate A, C, G, and T/U. For instance, for a 6-mer model, initially (iteration round 0), we assume the first base of the k-mer is the most significant base (Fig. 2 first plot titled *base shift:0*). Then we create a 1-mer table by ignoring the rest of the bases in the k-mer. There can only exist 4 different 1-mers (A, C, G, T/U), i.e., the current levels for AXXXXX and AYYYYY both now get mapped to 1-mer A. That is a many-to-one mapping (1024 current level mappings to 1-mer) per each 1-mer. There will be 4 current level

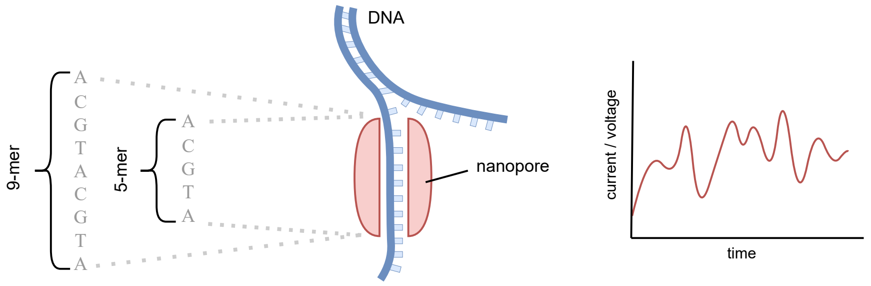


Figure 1: Depending on the size of the nanopore the k-mer inside the pore varies in length. In addition, depending on how the k-mer-model was generated, the most significant base may or may not be the base at the center of the k-mer.

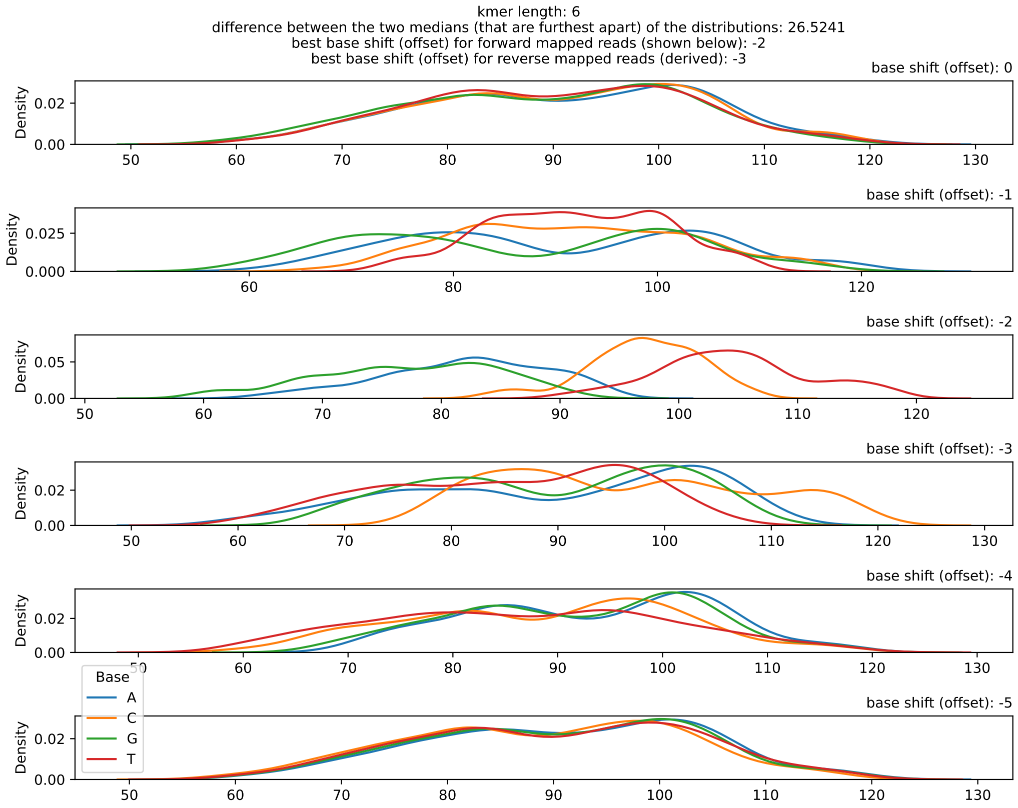


Figure 2: DNA R9.4.1 6-mer model

distributions (each distribution containing 1024 current levels) per base A, C, G, and T/U (Fig. 2 first plot titled *base shift (offset) :0*). The narrower the individual distributions are and the more the 4 distributions are separated from each other; the stronger our assumption becomes. In the next iteration (round 1), we assume the second base of the k-mer is the most significant base and repeat the experiment (Fig. 2 second plot titled *base shift (offset):-1*). Likewise, we do k iterations (Fig. 2). Then we choose the iteration where the 4 distributions had the best level of separation (third plot titled *base shift (offset):-2* in Fig. 2 that relates to iteration 2).

Table 1: K-mer Model specifications (*Nanopolish/f5c* models)

| **Chemistry** | **k-mer size** | **Most significant base index (0-based)** | **Figure No.** |
|---|---|---|---|
| DNA R9.4.1 | 6-mer | 2 | 2 |
| RNA R9.4.1 | 5-mer | 1 | 3 |
| DNA R10.4.1 | 9-mer | 6 | 4 |

In DNA models, looking at the current level distributions for each nucleotide of the most significant base index, we observe that the highest median current is for the nucleotide 'T' followed by the second highest median current for the nucleotide 'C'. The median currents for the nucleotide 'A' and 'G' are the lowest and they are close to each other.

To generate similar density plots for a new k-mer model, the following command can be used (more details in section 2).

```
squigualiser calculate_offsets --use_model --model ${MODEL_PATH} -o ${OUTPUT_PDF_PATH}
```

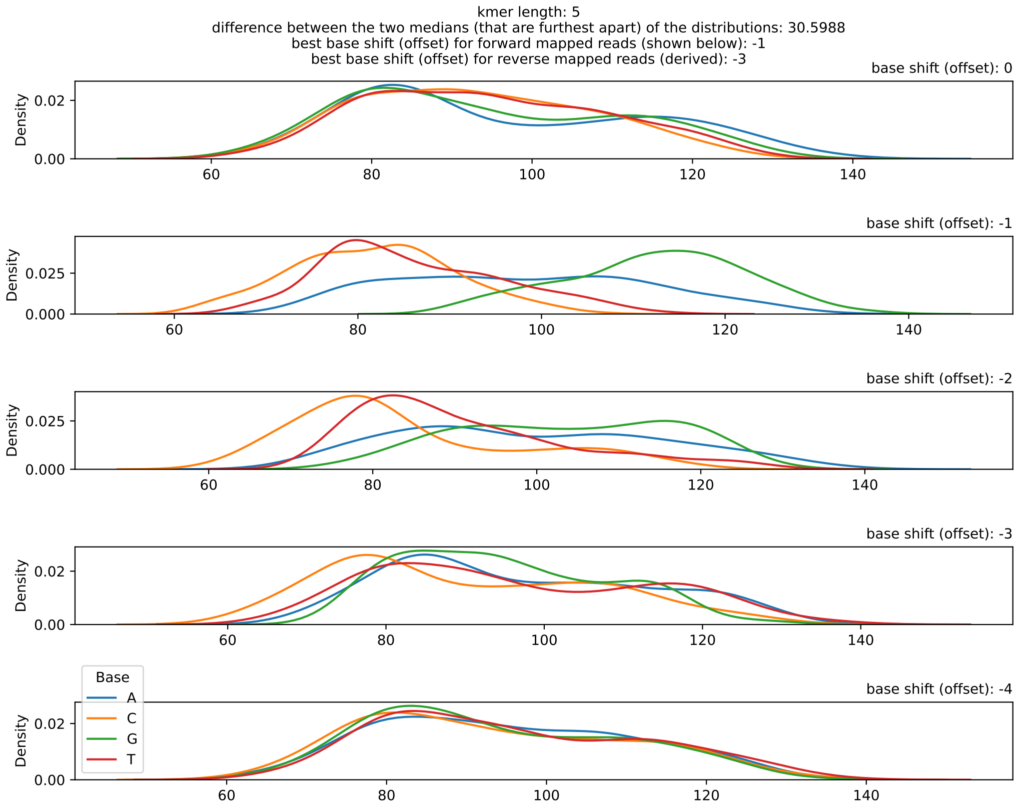


Figure 3: RNA R9.4.1 5-mer model

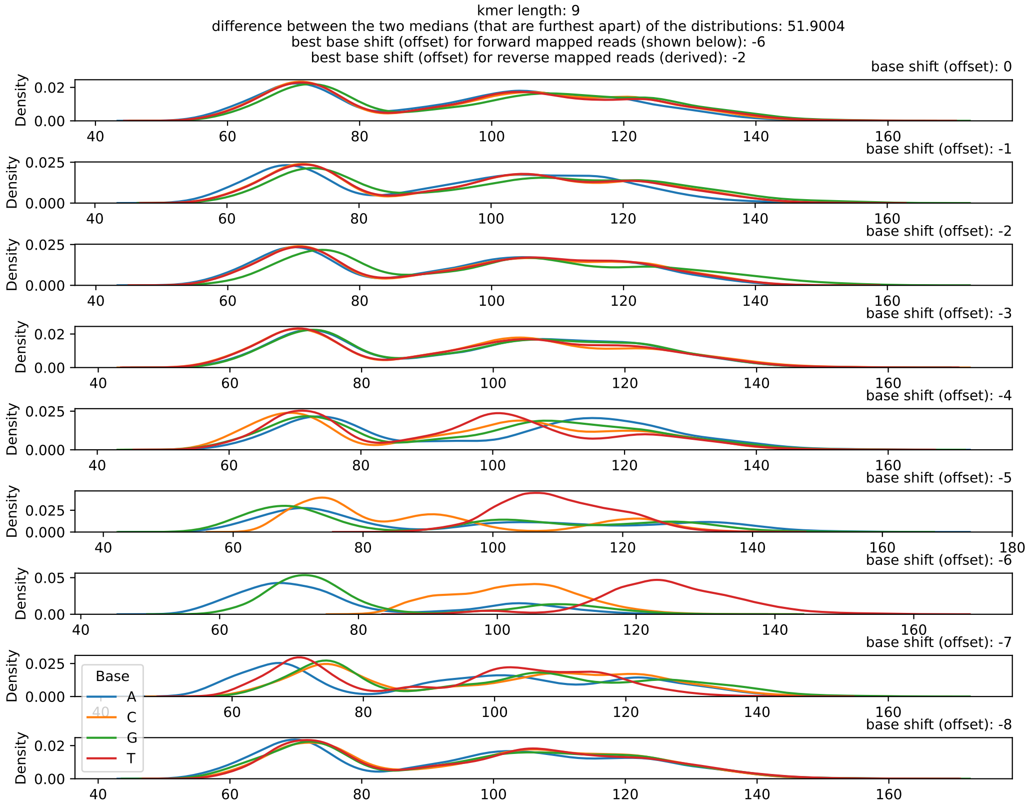


Figure 4: DNA R10.4.1 9-mer model

#### 2 *Squigualiser calculate_offsets* subtool

As the reader is now familiar with the concept of th most significant base, we will now delve into details of the *Squigualiser calculate_offsets* subtool. *calculate_offsets* can calculate the signal-to-most significant base alignment and has two modes. As mentioned in the main text, k-mer-to-base shift correction is necessary for both signal-to-kmer alignment (explained in *Mode 1*) and signal-to-base alignment (explained in *Mode 2*, **Supplementary Note 2**).

##### 2.1 Mode 1

*Squigualiser calcualte_offsets* when run in *Mode 1* setting will calculate the most significant base of a k-mer model. Subsequently, any signal-to-k-mer alignment that used the k-mer model should be transformed to signal-to-most significant base alignment. This transformation is done inside *Squigualiser*. *Nanopolish/f5c* and *Squigulator* default k-mer models are listed as profiles in Table 2. The user can provide the profile name when using *Squigualiser plot* or *plot_pileup* subtools. An example is given below.

```
squigualiser plot_pileup [OPTIONS] --profile profile_name -f genome.fasta -s reads.blow5
    -a eventalign.bam -o output_dir --region region
```

If the signal alignment method used a custom k-mer model then the user is advised to run *calcualte_offsets* to find the most signficant base index of the k-mers in the model as follows.

```
squigualiser calculate_offsets --use_model --model ${MODEL_PATH} -o ${OUTPUT_PDF_PATH}
```

Above command will output *base_shift* values for forward mapped reads and reverse mapped reads. The values can be used in *Squigualiser plot* or *plot_pileup* as follows.

```
squigualiser plot_pileup [OPTIONS] --base_shift forward_base_shift -f genome.fasta -s
    reads.blow5 -a eventalign.bam -o output_dir --region region
```

```
squigualiser plot_pileup [OPTIONS] --plot_reverse --base_shift reverse_base_shift -f
    genome.fasta -s reads.blow5 -a eventalign.bam -o output_dir --region region
```

The derivation of the *base shift* value for reverse mapped reads is illustrated in Fig. 5

Table 2: Model Parameters

| k-mer Model Profile | Offset (Forward) | Offset (Reverse) |
|---|---|---|
| kmer_model_dna_r9.4.1_450bps_5_mer | -2 | -2 |
| kmer_model_dna_r9.4.1_450bps_6_mer | -2 | -3 |
| kmer_model_rna_r9.4.1_70bps_5_mer | -3 | -1 |
| kmer_model_dna_r10.4.1_e8.2_400bps_9_mer | -6 | -2 |

The pseudo code given below explains the computation done inside *calculate_offsets Mode 1*.

```
find_most_significant_base(kmer_model){

    for base_index in kmer_length{
        for kmer, current_level in kmer_model{
            bin current_level to one of the four bins A,C,G and T
        }

        draw four density distributions, one for each bin
        calculate difference (diff) between highest and lowest medians of the
    distributions
    }

    select the maximum difference (max_diff) among the differences (diff)
    pick the corresponding base index (base_index) as the most significant base index

    return  max_diff, base_index
}
```

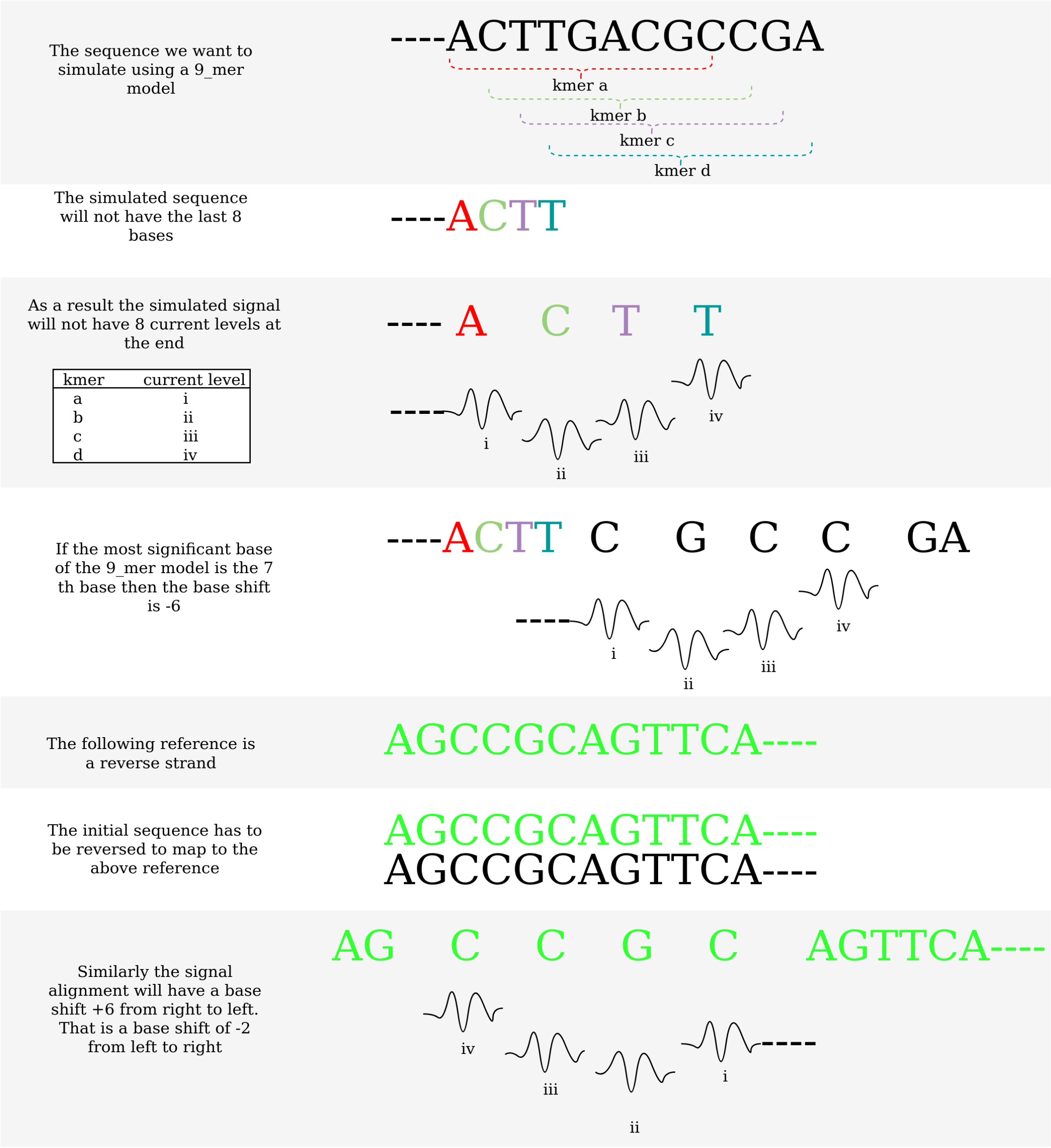


Figure 5: Illustration of deriving correct *base shift* value for reverse mapped sequences

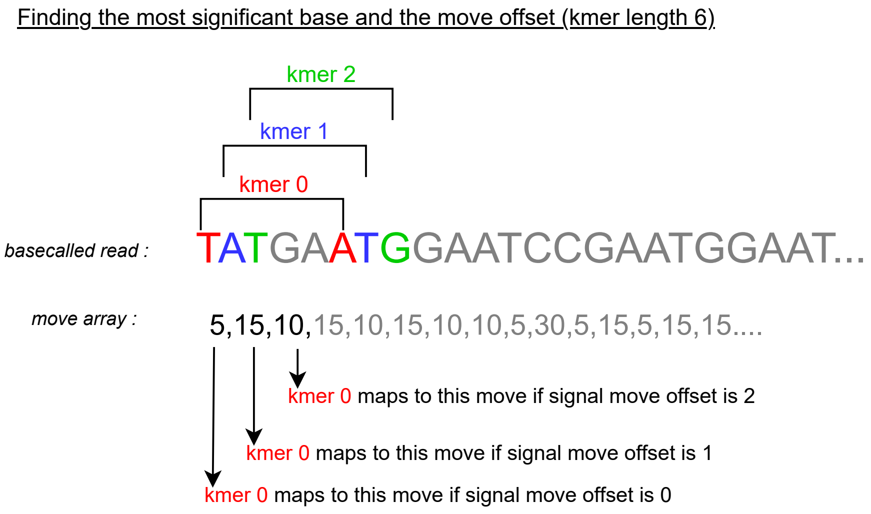


Figure 6: Illustration of how *squigualiser calculate_offsets* Mode 2 finds the best *kmer_length* and *sig_move_offset* to generate the signal-to-most significant base alignment.

#### 2.2 Mode 2

As explained above and in **Supplementary Note 2**, even though basecaller's move table is a signal-to-base alignment, depending on the basecalling model it might not be the signal-to-most significant base alignment. *Squigualiser calcualte_offsets* when run in *Mode 2* setting will output *kmer_length* and *sig_move_offset* values to be used in *Squigualiser reform* subtool.

```
squigualiser calculate_offsets -p input.paf -s reads.blow5 -f reads.fastq
```

```
squigualiser reform [OPTIONS] -k kmer_length -m sig_move_offset -c --bam movetable.bam -o
     reform.paf
```

The following pseudo code and Fig. 6 should help in understanding how the most significant base index is determined in Mode 2.

```
DEFAULT_KMER_LENGTH = 6
best_m = 0
best_max_diff = -infinity
most_significant_base_index = 0
recommended_sig_move_offset = 0
for m in range(0,DEFAULT_KMER_LENGTH):
 # create a kmer model by iterating the fastq sequence, assigning each kmer the current
    value pointed by the move and pass it to Mode 1
    local_max_diff, base_idx = finding_the_most_significant_base(kmer model)
    if best_max_diff < local_max_diff:
        best_max_diff = local_max_diff
        best_m = m
        most_significant_base_offset = base_idx
        if most_significant_base_offset == 0:
            recommended_sig_move_offset = m
# assumption: the recommended_sig_move_offset is the m value where the
    most_significant_base_offset is zero and as we increment m by 1
    most_significant_base_offset also gets incremented by 1)

# confirm that the assumption is valid by checking...
if recommended_sig_move_offset == best_m - most_significant_base_offset
    then recommended_kmer_length = recommended_sig_move_offset + 1
# else increment the DEFAULT_KMER_LENGTH by 1 and further analyse the density plots
    generated.
```

Mode 2 can optionally take a read id to plot a density plot (similar to Fig. 2, 3, and 4) for the particular signal-to-read alignment as follows,

```
squigualiser calculate_offsets -p input.paf -s reads.blow5 -f reads.fastq -o out.pdf --
    read_id READ_ID
```

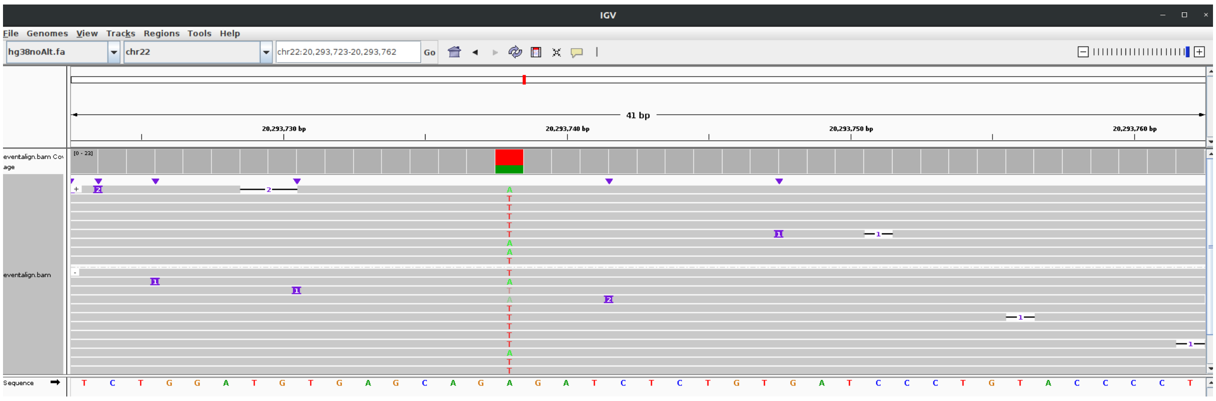


Figure 7: IGV reference-to-read pileup with heterozygous A/T SNV at chr22:20,293,723-20,293,762
.

### 3 Validating the k-mer-to-base-shift correction using an example

#### 3.1 Preprocessing

In this example let's look at a heterozygous A/T SNV found in the humangenome (hg38). This example only focus on forward-mapped reads. At site chr22:20,293,738 there is A (Read 1,7, and 8) and T (Read 2,3,4,5,6, and 9)(Fig. 7).

Fig.8 shows three signal alignments. The first is the signal-to-reference alignment using *f5c eventalign*. The second and the third are the ideal simulated signals using the reference with and without the SNV respectively. Grey circles mark the SNV we are interested in. As shown in the simulated signals there must be a clear jump up in the signals at A21 when it is a T (third track). However, the *eventalign* algorithm has failed to properly align it the specific position for the reads that have the SNV. It has introduced deletions, and aligned the jump before (at A19) or after (at T24). This phenomenon is not correct but expected as the reads were aligned to the original reference where an A was present, not a T. To rectify this we aligned the reads again to a new reference where the particular A was replaced with a T. Then, we executed *eventalign* with the new read alignment file and the new reference. Subsequently, we selected the signal alignment records from the correct *eventalign* outputs (Read 1,7,8 and 2,3,4,5,6,9 extracted from the original and new outputs respectively). Then we merged the signal alignment records to create a single signal alignment file. It was used to generate a better signal alignment pileup (Fig. 9). This pre-processing workflow is summarised in Fig. 10.

#### 3.2 Statistical evaluation

This section is an extension of the kmer-to-base shift correction results presented in Fig. 3 of the main text. We have chosen 250 heterozygous A/T variants in chromosome 22 based on HG001 GRCh38 GIAB high-confindence variants dataset. We plotted the normalised signal values at the heterozygous SNV for different base offsets, i.e., if the k-mer length is 9, then offset 0 means the signal event aligns with the 0th base of the k-mer, and offset -6 means that the signal event aligns with 6th base of the k-mer. Fig. 11 shows seven more signal value distribution plots for seven SNV positions (first seven columns) including the collective distribution for 250 SNVs (last column). We observed that when the the offset is -6; we get the highest degree of bimodality (the SNV causes the signal to have a different current level at the SNV position). At single SNV resolution this observation is more pronounced.

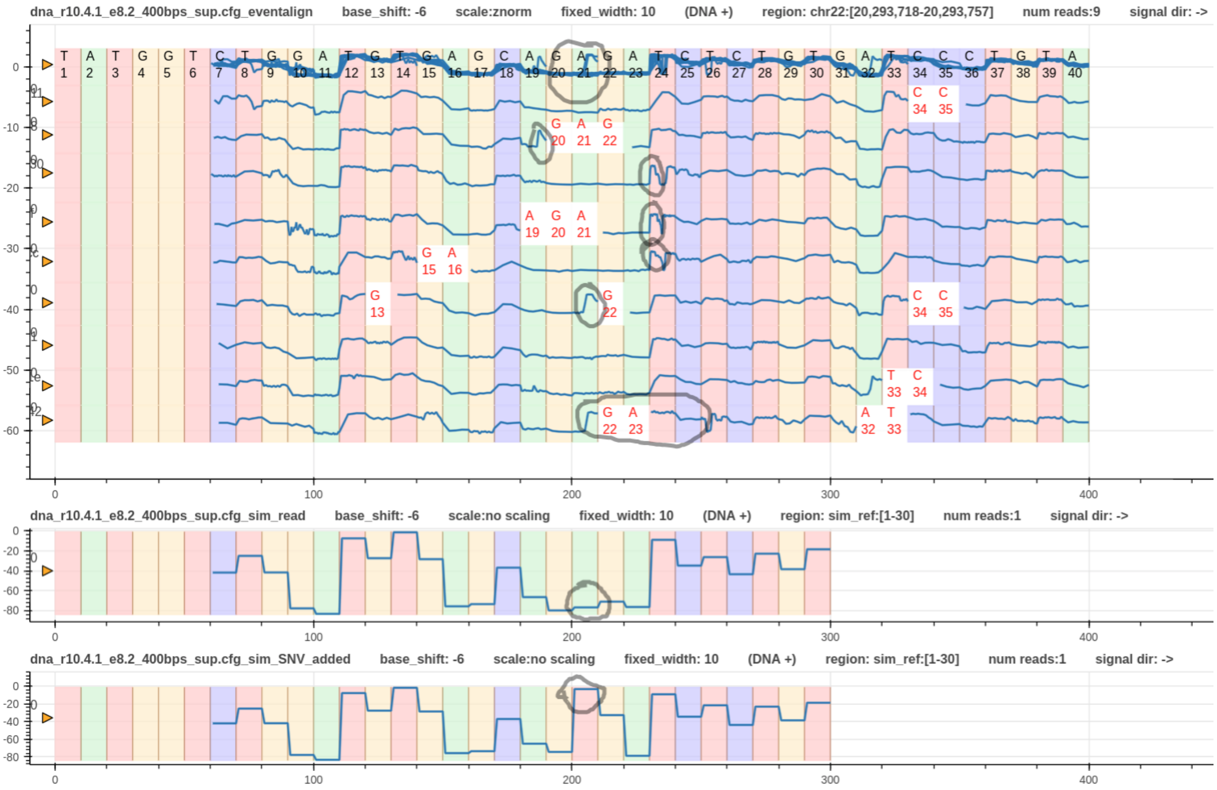


Figure 8: First signal alignment track is the *f5c eventalign*. The second and third tracks show the simulated (expected) ideal signal shape using *squigulator* with and without the SNV, respectively

.

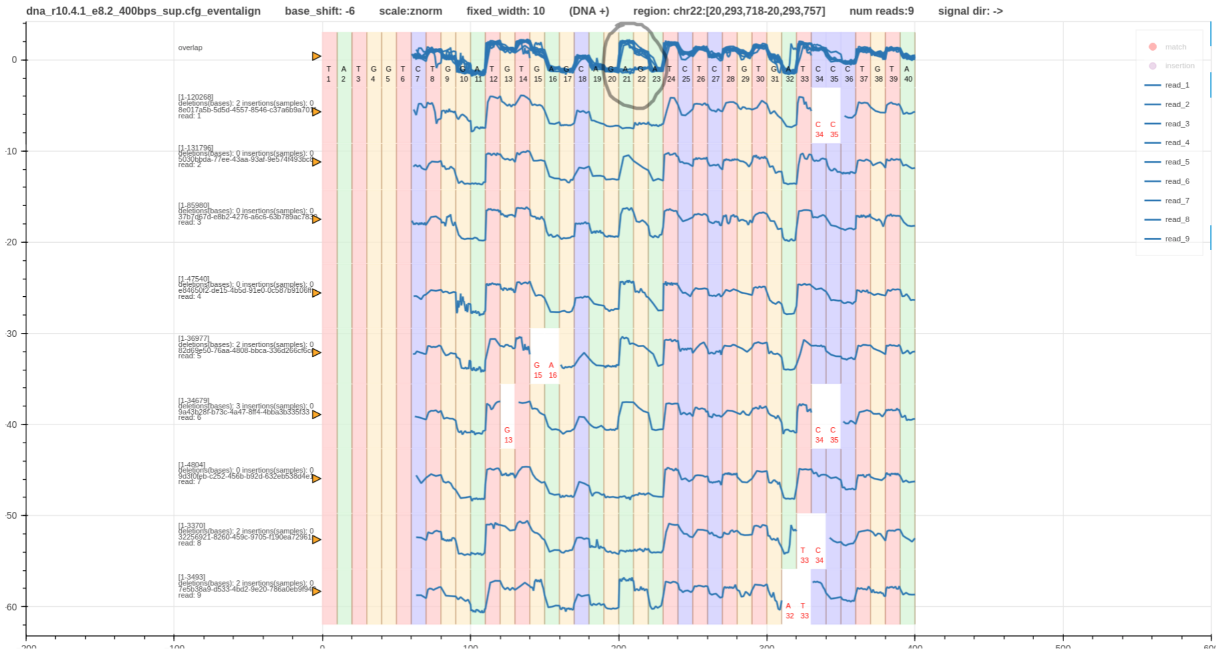


Figure 9: The signal aligment pileup after aligning the signals to the correct reference, i.e., the signals with the SNV were aligned to the reference with the SNV

.

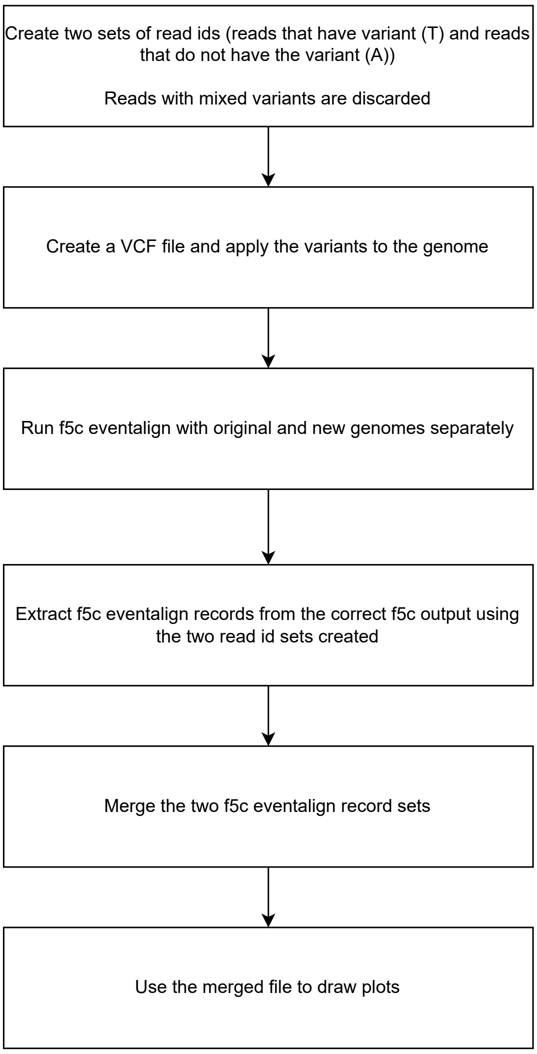


Figure 10: The preprossessing workflow to generate more sensible *eventalign* signal alignment plots
.

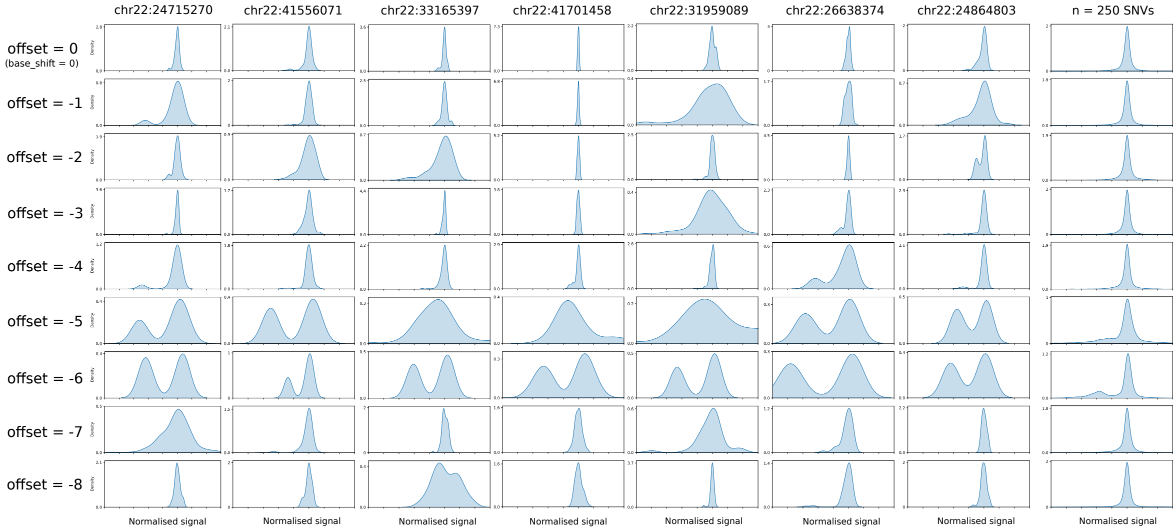


Figure 11: The distribution of normalised signal values aligned at the SNV position with different offsets. The first 7 columns are individual distributions and the last column is the collective distribution of 250 SNVs.

.

### Supplementary Note 4: Visualising a genomic region that has a CpG methylated site

Hiruna Samarakoon, Kisaru Liyanage, James M. Ferguson, Sri Parameswaran, Hasindu Gamaarachchi, Ira W. Deveson

February 16, 2024

Methylated cytosine bases cause differences in the nanopore current level compared to non-methylated bases. This document explains how such current differences can be observed using a squigualiser pileup plot. Assume we already have methylation frequency information calculated based on the output of *f5c* call-methylation command. The steps to visualise a region with a half-methylated CpG site that effectively demonstrates differences in current levels are given below.

1. Filter a genomic site from the methylation frequency file where the frequency is approximately 0.5.
2. Filter the read ids from the call-methylation TSV file where the log likelihood ratio is less than $-2$ and above $2$ for unmethylated and methylated reads, respectively.
3. Use the filtered read ids to create a pileup plot:

```
squigualiser plot_pileup [OPTIONS] -l readlist -f reference.fa -s reads.blow5 -a
    eventalign.bam --region region
```

Let's now examine an example from our HG002 DNA 10.4.1 dataset (SRA: SRR23215366). In this example, we considered the + strand reads covering the site chr1:92790687 (1-based coordinate) with a methylation frequency of 0.542. As shown in the overlap plot in Fig.1 (top track), the site C38 (marked by red block) has two distinct current levels. The higher current level corresponded with the unmethylated read_ids in the methylation calls table (Table 1). The lower current level corresponded with the methylated read_ids. Out of the 8 total reads in the pileup, 3 were unmethylated, and 5 were methylated.

Table 1: Methylation calls from *f5c* call-methylation. Note that positions have been manually edited to be 1-based to match with the coordinate convention of squigualiser.

| Chromosome | Strand | Position | Read Name | Log Lkhd Ratio | Log Lkhd Methylated | Log Lkhd Unmethylated | No. of Calling Strands | No. of Motifs |
|---|---|---|---|---|---|---|---|---|
| chr1 | + | 92790687 | d2d2c018-f49f-436b-a038-e441f95a49ad | -2.69 | -162.08 | -159.39 | 1 | 1 |
| chr1 | + | 92790687 | 5ae057b3-cde0-4248-a905-d18a75079ba2 | 4.6 | -127.22 | -131.82 | 1 | 1 |
| chr1 | + | 92790687 | caad6a89-8ad5-485d-9ed9-903e99ed1b6a | 3.8 | -196.16 | -199.95 | 1 | 1 |
| chr1 | + | 92790687 | 0aa2c317-d8c4-4861-ac6f-87bab7fff2d8 | 4.77 | -134.04 | -138.81 | 1 | 1 |
| chr1 | + | 92790687 | fdb21051-2388-446d-acf2-a6848669ced8 | 2.62 | -84.27 | -86.89 | 1 | 1 |
| chr1 | + | 92790687 | 1e12424f-dcab-4cd0-83c3-79afc1326199 | -4.41 | -120.27 | -115.86 | 1 | 1 |
| chr1 | + | 92790687 | 362dc957-024e-4c56-a26a-c7fe4cd07304 | 4.15 | -198.4 | -202.56 | 1 | 1 |
| chr1 | + | 92790687 | 6b960395-8f61-4b8f-880d-e1852a5556a8 | -2.78 | -147.48 | -144.7 | 1 | 1 |

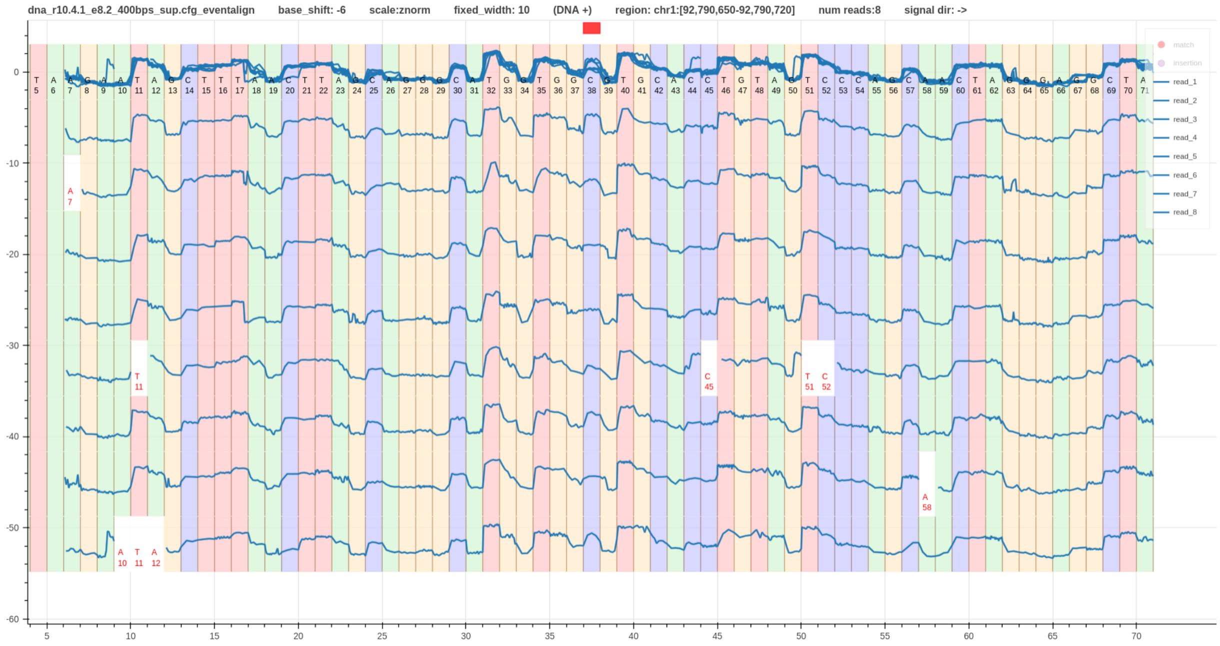


Figure 1: *Squigualiser pileup* plot including the overlap of the signals at the top. The methylation detection conducted using in *f5c* indicated that the site C38 has a methylation frequency of 0.542.

### Supplementary Note 5: Visualising RNA modifications

Hiruna Samarakoon, Kisaru Liyanage, James M. Ferguson, Sri Parameswaran,
Hasindu Gamaarachchi, Ira W. Deveson

February 17, 2024

In this document, we will look at synthetic direct RNA data from ONT rna_r9.4.1_70bps chemistry. We downloaded the publicly available data (Table 1) comprising of positive and negative controls of pseudouridine modifications.

Table 1: Dataset Information

| **Dataset name** | **SRA accession description** | **Preparation** |
|---|---|---|
| unmodified (UnMod_37C) | direct RNA-seq of GFP mRNA (SRR22888949) | synthetic RNA with all sites unmodified |
| modifiedU (Mod_37C) | direct RNA-seq of GFP mRNA (SRR22888950) | synthetic RNA with all Us modified |

The following steps were executed.

1. Basecall the two datasets using guppy_v6.3.7 with the model rna_r9.4.1_70bps_hac_prom.cfg.
2. Align the reads to the synthetic reference using *minimap2*.
3. Generate a signal-to-reference alignment using *squigualiser realign* program (see **Supplementary Note 2**).
4. Generate another signal-to-reference alignment using *Nanopolish/f5c eventalign*.
5. Evaluate signal-to-reference alignments by visualising the signal pileups (using *Squigualiser plot_pileup*).
6. Improve *Nanopolish/f5c* signal-to-reference alignment by training the k-mer model using *Nanopolish train* program.
7. Evaluate the efficiacy of *Nanopolish training* by observing the signal-to-reference alignments generated using the k-mer models after each training round.
8. Observe the feature differences in the unmodified and modified signals.

When using the default basecalling parameters, the reported pass read percentages were 92% for the unmodified dataset and 62% for the modifiedU dataset. The very low pass percentage (62%) for the modifiedU dataset indicates that the synthetically modified U has very different signal features than the normal U. This effect has made the basecalling models erroneous around U bases.

This can be further confirmed by looking at the metrics after aligning the passed reads to the synthetic reference (Table 2). The modifiedU dataset's mapping percentage is very low (53%) compared to the unmodified dataset's mapping percentage (99%). The k-mer size and alignment score parameters were changed in minimap2 to get a better mapping percentage (97%, Table 2).

By comparing the reference-read alignments on IGV, we can observe that the basecaller has basecalled almost all modifiedUs as C (Fig. 1 & 2).

Table 2: *Minimap2* alignment statistics

| | **minimap2 params** | | **Reads** | | | | **Bases** | | | | |
|---|---|---|---|---|---|---|---|---|---|---|---|
| **Dataset** | **-K (k-mer size)** | **-s (DP score)** | **Total** | **Mapped** | **Unmapped** | **Mapped %** | **Total** | **Mapped** | **Mapped %** | **Mismatches** | **Error rate** |
| unmodified | default (15) | default (80) | 4622 | 4615 | 7 | 0.998 | 5137565 | 4922590 | 0.959 | 631666 | 0.129 |
| modifiedU | default (15) | default (80) | 445310 | 238402 | 206908 | 0.5358 | 459273108 | 197679705 | 0.431 | 54221392 | 0.275 |
| modifiedU | 10 | 50 | 445310 | 432604 | 12706 | 0.9718 | 459273108 | 347414498 | 0.757 | 97745894 | 0.282 |

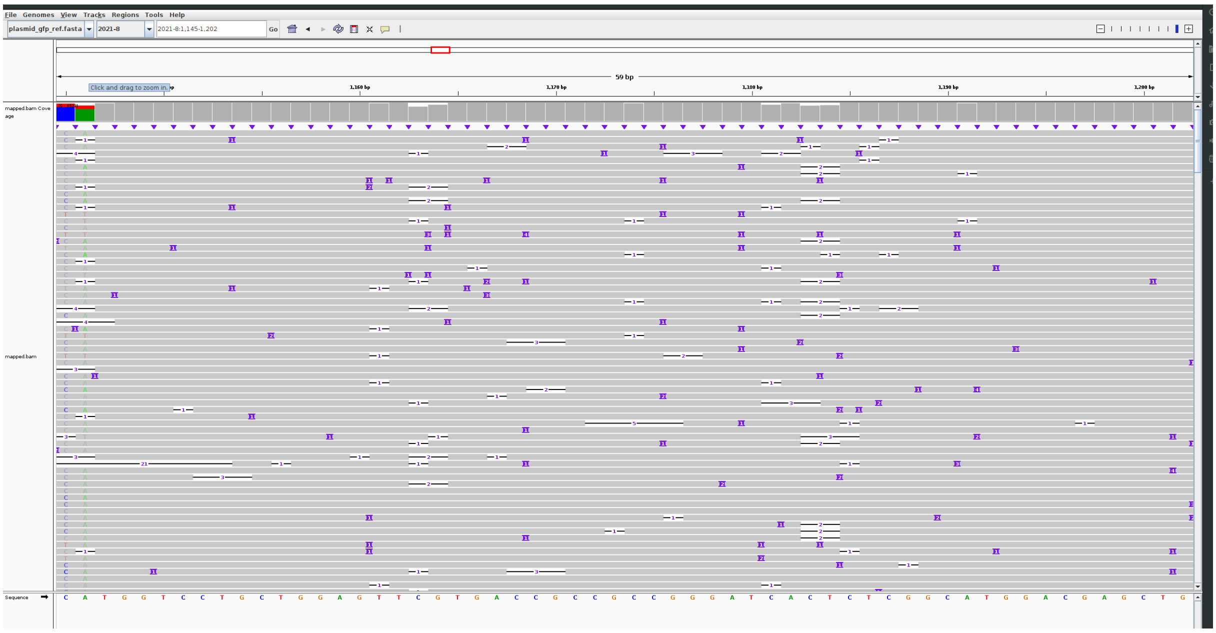

Figure 1: Unmodified read-to-reference alignment

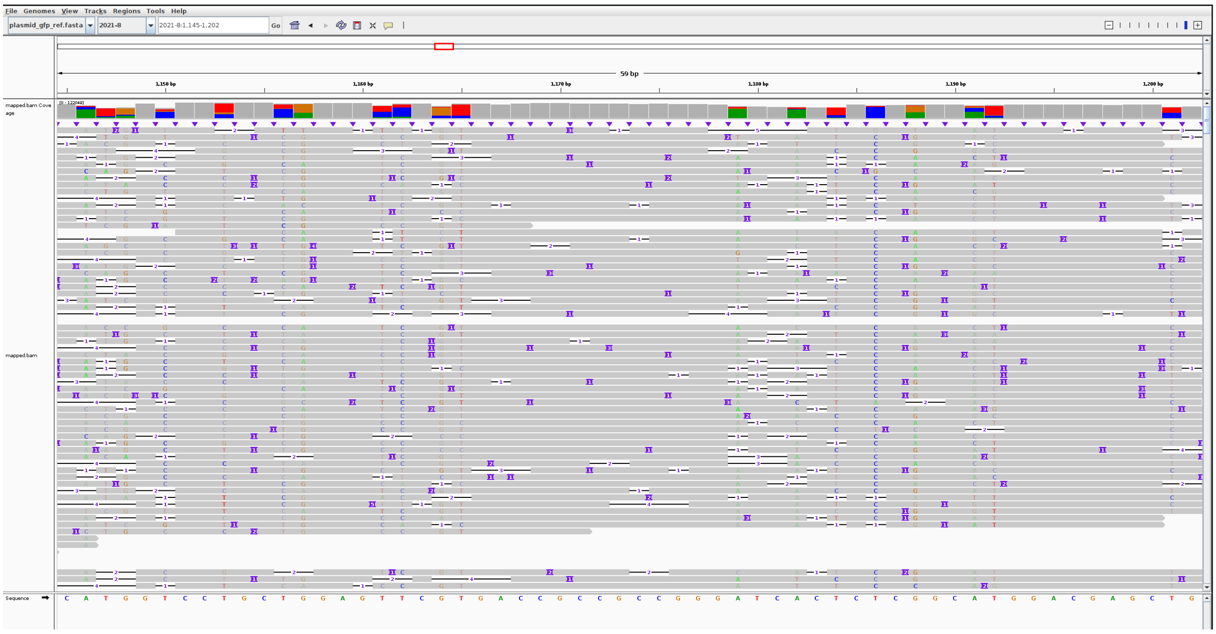

Figure 2: ModifiedU read-to-reference alignment

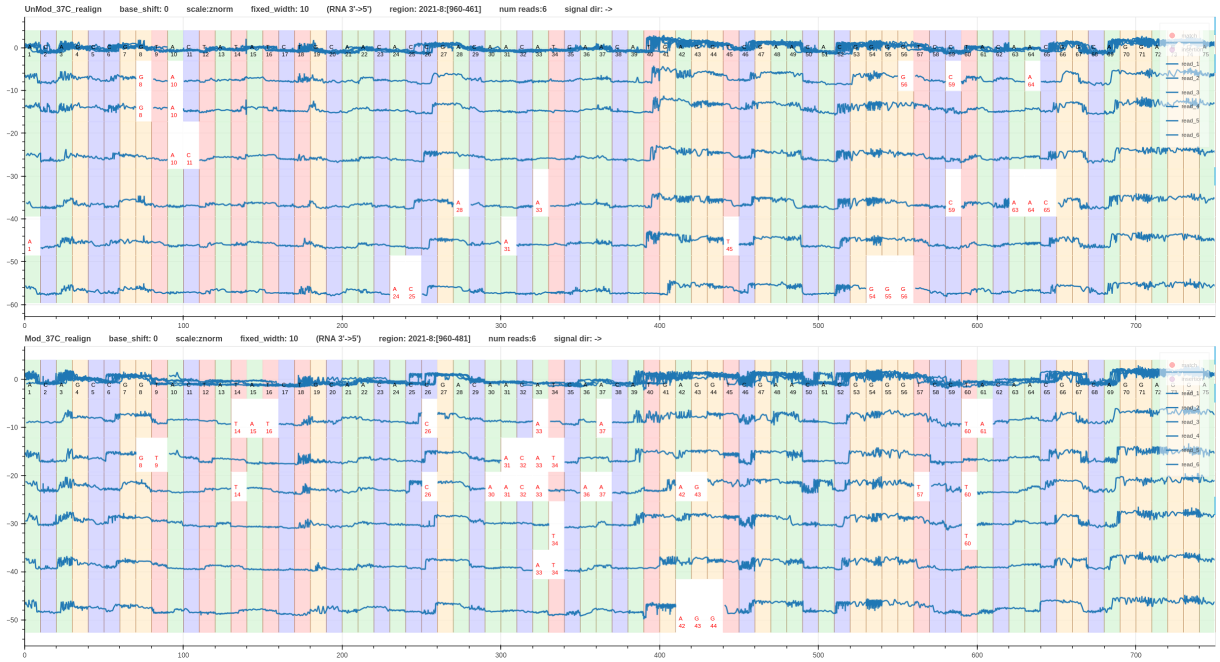


Figure 3: Unmodified and modifiedU signal-to-reference alignment using *Squigualiser realign*

Now let's proceed to the signal alignment stage.

*Squigualiser realign* method appear to be robust around modified Us as it uses the basecaller's move table (it does not matter if it is a C or U as long as it is a counted as a move). However, the boundaries of signal alignment are not refined in this method (Fig. 3).

Fig. 4 shows a comparison between the unmodified and modifiedU data signal alignment using *Nanopolish/f5c eventalign*. The signal alignment is erroneous around U bases in the modifiedU data. K-mers with modified Us have different current levels than the k-mers with unmodified Us. Since *Nanopolish/f5c eventalign* relies on k-mer model, it is critical that the k-mer model has the updated current levels for the modified k-mers. Therefore, *Nanopolish train* program was used to train a k-mer model for the modified data.

The *Nanopolish* training was carried out for 10 rounds. The trained k-model at each round was given as a custom model to *f5c* eventalign program to align the modifiedU signal data. However, it was observed that the signal alignment accuracy did not improve after the first two rounds (Fig. 5). Hence, the k-model trained in round 1 (0-based) was used to align the modifiedU data (Fig. 6).

```
f5c eventalign -b minimap2.bam -r reads.fastq -f genome.fa -s reads.blow5 -a eventalign.
    bam --kmer-model roundX.model
```

Fig. 7 takes a closer look at the current level differences of the k-mers with U between unmodified and modifiedeU datasets. We observe that the current level of a k-mers with modified U drops below the current level of the corresponding unmodified k-mer. This observation is more pronounced when there are several modified Us together.

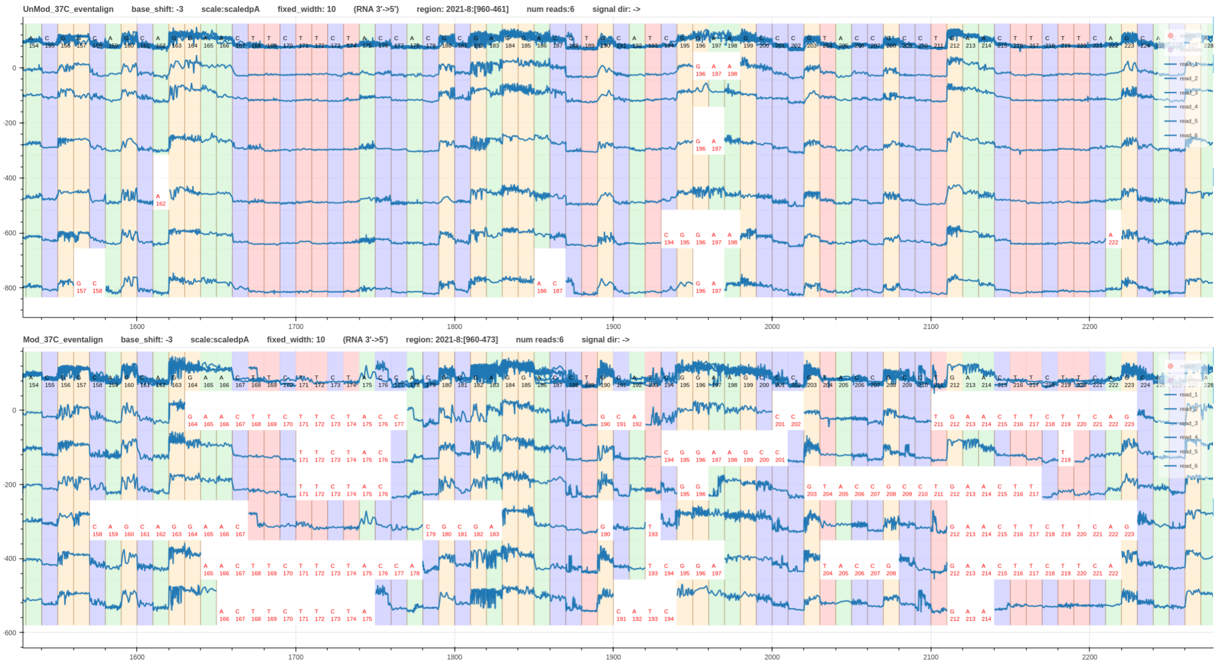

Figure 4: Unmodified (top pileup track) and modifiedU (bottom pileup track) signal-to-reference alignment using *Nanopolish/f5c eventalign*

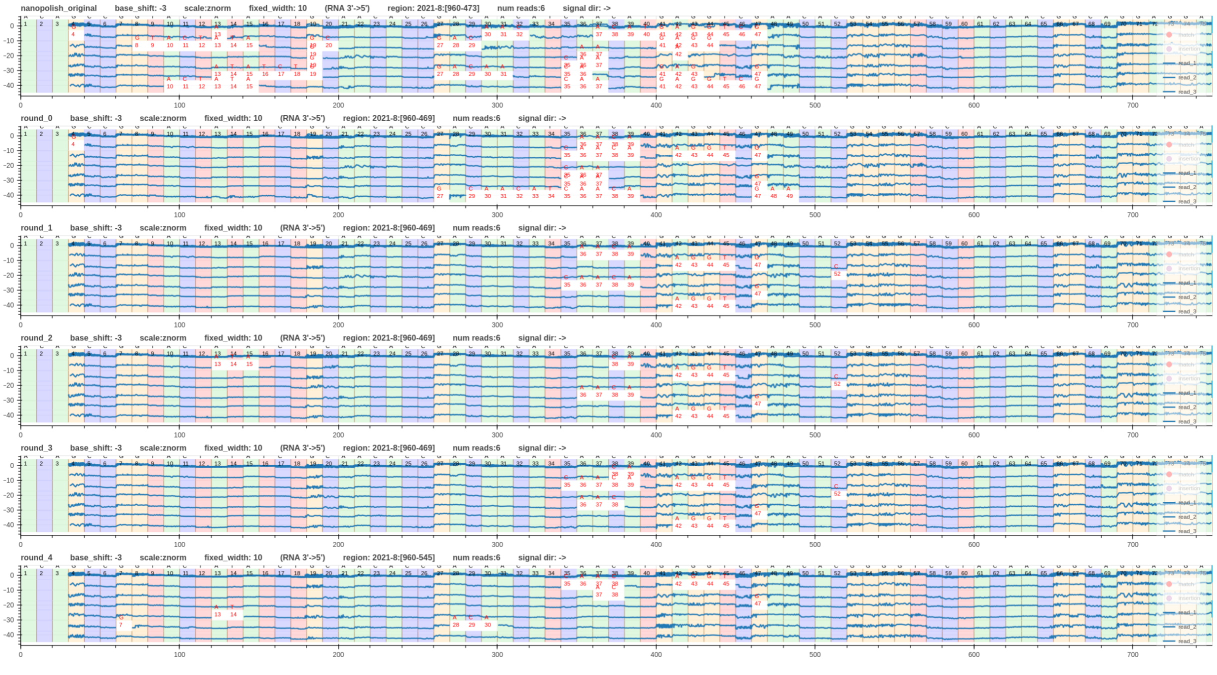

Figure 5: ModifiedU signal-to-reference alignment using *Nanopolish/f5c eventalign* with trained k-mer models. The topmost pileup track is for the original k-mer model and the subsequent tracks are for trained models from training rounds 0 to 4.

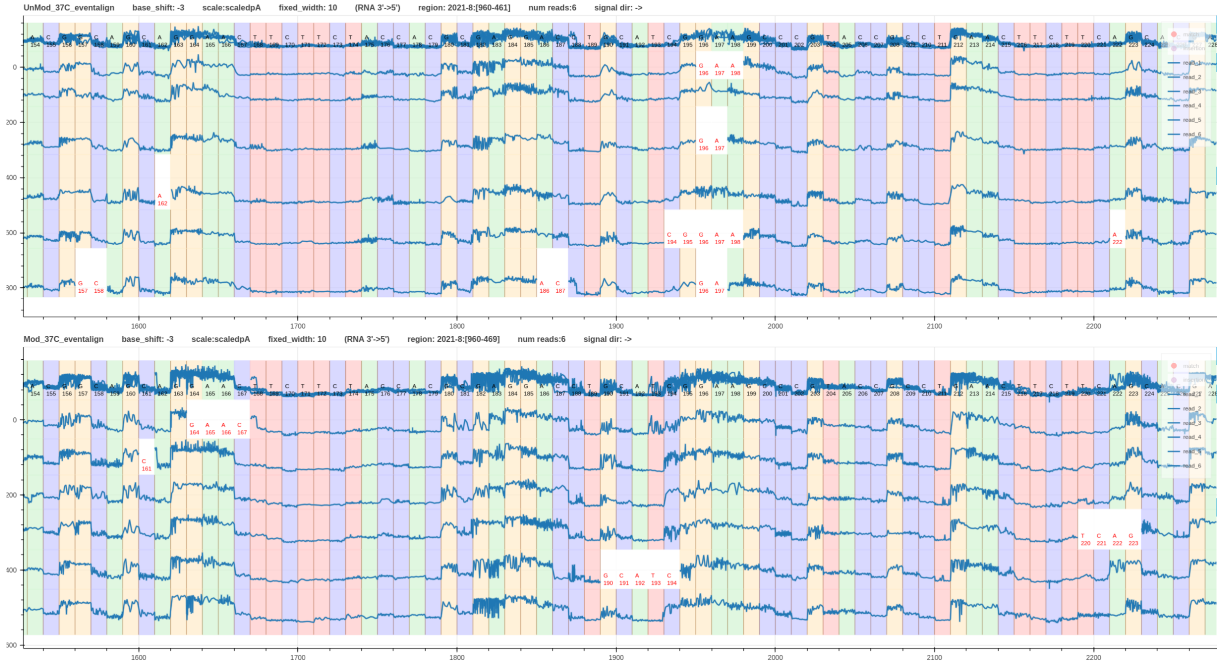


Figure 6: Unmodified signal-to-reference alignment with original k-mer model (top pileup track) and modifiedU signal-to-reference alignment with trained k-mer model (bottom pileup plot). Alignment performed using *Nanopolish/f5c eventalign*.

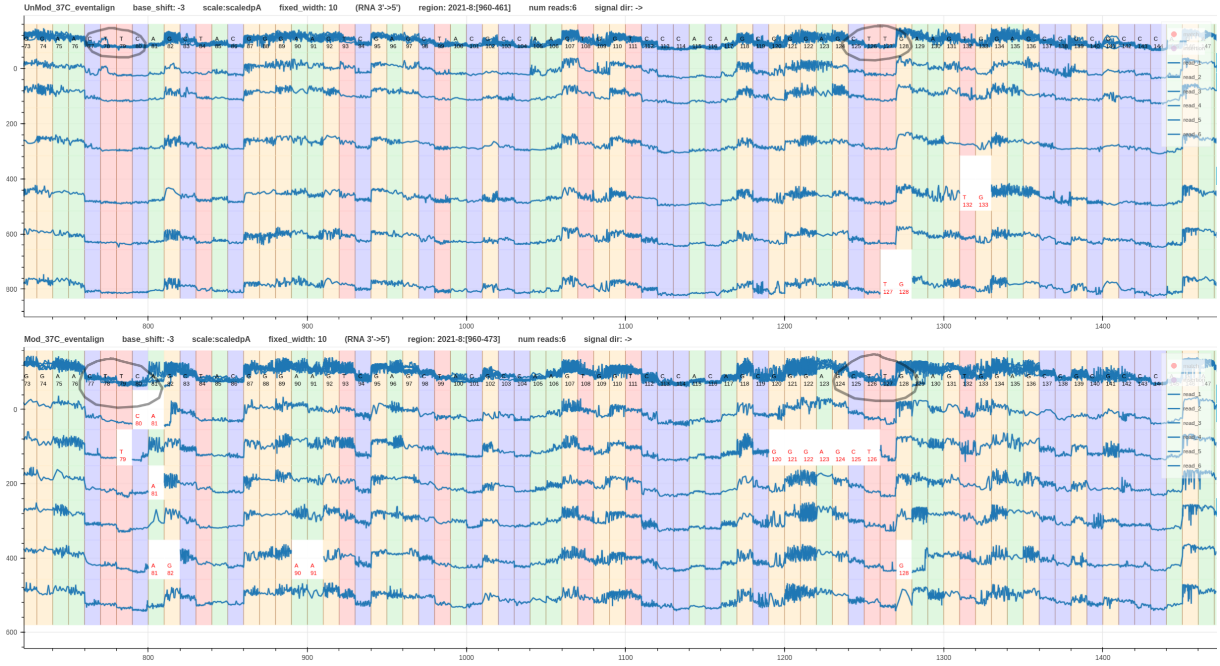


Figure 7: Unmodified and modifiedU current level comparison

### Supplementary Note 6: Squigualiser plot conventions

Hiruna Samarakoon, Kisaru Liyanage, James M. Ferguson, Sri Parameswaran,
Hasindu Gamaarachchi, Ira W. Deveson

February 19, 2024

For the best user experience *Squigualiser* conforms to the plot conventions that are described in Fig. 1.

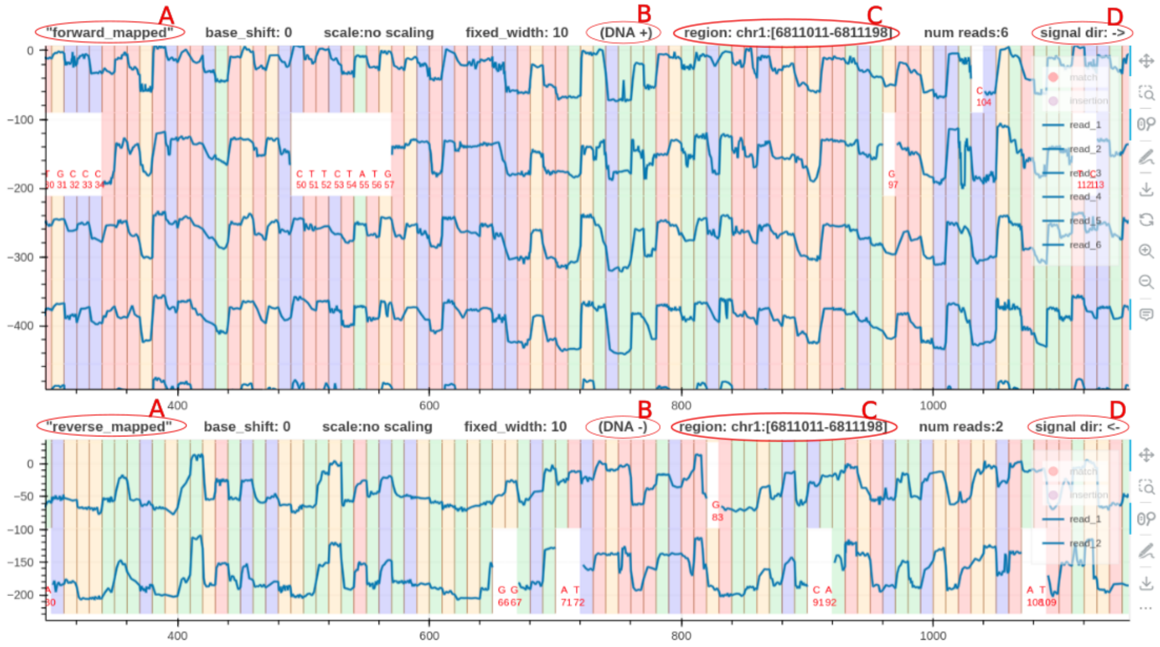


Figure 1: Plot conventions of squigualiser
**A** is a descriptive tag name to identify the plot.
**B** indicates whether the positive or negative strand was used as the reference to align the signals. For RNA this will be RNA 3'->5'. *Squigualiser* only supports RNA reads mapped to the transcriptome.
**C** always indicates the region using the positive strand coordinates, regardless of the forward and reverse mapped plots.
**D** indicates the true sequencing direction of the signals.

### Supplementary Note 7: Squigualiser plot annotations

Hiruna Samarakoon, Kisaru Liyanage, James M. Ferguson, Sri Parameswaran,
Hasindu Gamaarachchi, Ira W. Deveson

February 19, 2024

*Squigualiser plot* and *plot_pileup* can take a BED file that has annotation information. The BED format can store color values and BED line names in addition to the genomic positions. The BED file used in generating the signal pileup shown in Fig. 1 had three BED lines (*POS1, POS2, and POS3*) and hence, the plot has three BED lines.

```
squigualiser plot_pileup [OPTIONS] --bed annotation.bed --print_bed_labels -f reference.
    fa -s reads.blow5 -a eventalign.bam --region region
```

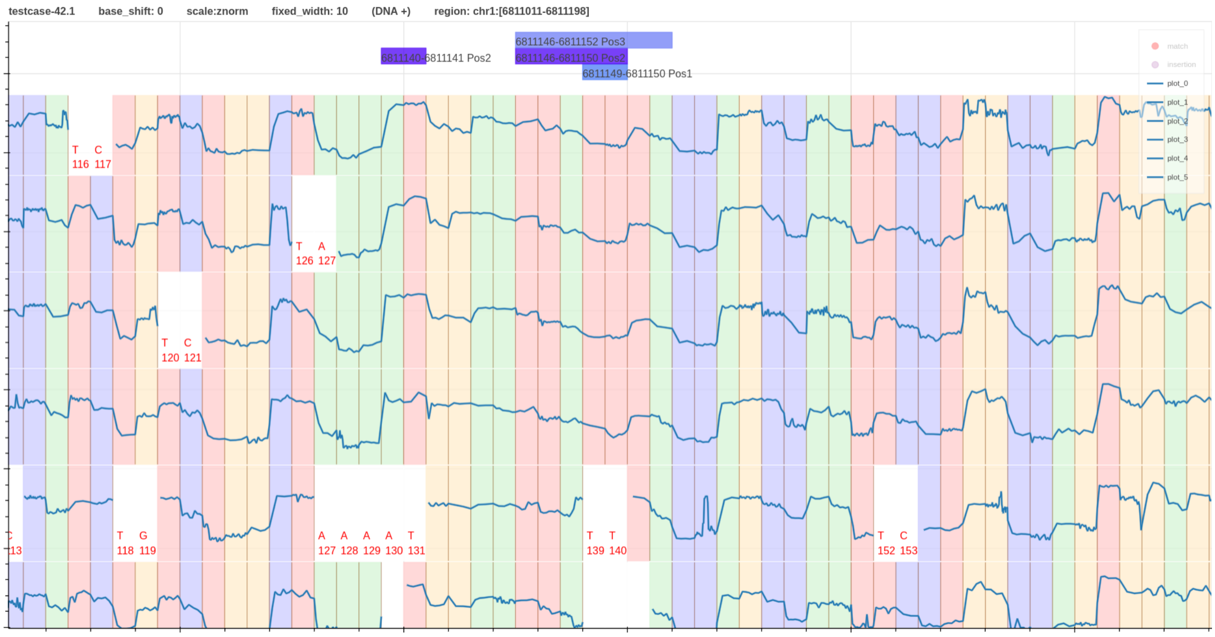


Figure 1: DNA signal pileup with three layers of annotation loaded using a single BED file.
